# HalluCodon enables species-specific codon optimization using multimodal language models

**DOI:** 10.64898/2026.03.31.715573

**Authors:** Yuxuan Lou, Shiya Mao, Tianhao Wu, Fan Xia, Zhaojie Zhang, Yisu Tian, Yu Li, Qian Cheng, Jun Yan, Xiangfeng Wang

## Abstract

Codon optimization is widely used in transgenic crop development, plant synthetic biology, and molecular farming to improve heterologous protein expression in plant cells. Increasing availability of plant omics data now enables optimization strategies that account for species-specific sequence features. We developed HalluCodon, a customizable framework that uses multimodal language models to design coding sequences tailored to individual plant species. The framework allows users to fine tune pre-trained protein and RNA language models with their own datasets to build species-specific codon optimization models. The current implementation includes base models trained on coding sequences and proteomes from fifteen plant species. HalluCodon generates coding sequences through a hallucination-based design strategy guided by two predictive modules that evaluate coding sequence naturalness (CodonNAT) and expression potential (CodonEXP). Benchmark tests using representative proteins show that the generated sequences reproduce host-specific codon usage patterns and support high expression levels in plant systems.

## Introduction

Codons are the basic units of the genetic code that determine the identity and order of amino acids during translation of genetic information from DNA into proteins in cells^1^. Because of codon degeneracy, multiple codons can encode the same amino acid. These codons are referred to as synonymous codons, meaning that a single protein sequence can be encoded by many different coding sequences (CDSs)^2^. However, synonymous codons are not used with equal frequency. Codon usage shows clear preferences that vary among species, among genes within the same genome, and even across different regions of a single gene^3, 4^. The choice of synonymous codons can strongly influence protein production through several mechanisms, including differences in tRNA abundance, effects on mRNA secondary structure, and the presence of sequence motifs such as restriction enzyme sites^5–7^. Designing coding sequences that match host codon usage preferences is therefore an important step in achieving efficient heterologous protein expression. This strategy is commonly referred to as codon optimization and is widely used in biotechnology. In plant systems, it is particularly relevant for plant synthetic biology and molecular farming, where plant cells are used as production platforms for natural or engineered proteins^8, 9^.

Many traditional codon optimization strategies rely on codon usage statistics derived from the host genome. One commonly used metric is the codon adaptation index (CAI), which measures how closely a designed coding sequence follows the codon usage preferences of the host species^10, 11^. However, codon frequency alone does not fully explain translation efficiency. In many genes, rare codons serve regulatory roles during translation. Their presence can locally slow ribosome movement and help ensure correct folding of nascent polypeptides. This effect is particularly important in regions encoding structurally complex protein domains. Consequently, replacing all rare codons with highly frequent ones may disrupt the natural translation dynamics of the protein, potentially leading to misfolding or aggregation of dysfunctional proteins^12^. In addition, excessive use of high frequency codons may cause imbalanced utilization of the cellular tRNA pool and negatively affect mRNA stability^6, 13^.

Together, these findings indicate that codon context is often more important than codon frequency alone. Codon usage patterns may therefore vary among genes within the same genome, reflecting gene specific regulatory constraints. Recent advances in deep learning have enabled new approaches to codon optimization by allowing models to learn complex sequence patterns directly from genomic data. Several Transformer based methods have been proposed for this task, including CodonTransformer, DeepCodon, and GEMORNA^14–16^. These tools demonstrate improved performance compared with traditional codon usage based approaches. However, most existing models are trained from scratch in an unsupervised manner and do not leverage pre-trained biological language models or experimental protein expression data. As a result, large amounts of coding sequence data are required for training. Applying these models to species outside the original training set therefore requires substantial data collection and retraining.

In mammals, codon optimization is frequently used in the design of mRNA vaccines, where minimizing innate immune responses to exogenous RNA sequences is a major consideration^17^. Endogenous mammalian mRNAs avoid immune activation through multiple mechanisms^18^, whereas synthetic mRNA sequences often require chemical modifications to reduce immunogenicity^19^. Codon optimized sequences must therefore satisfy both codon usage constraints and compatibility with these modifications. In contrast, plants generally do not mount immune responses against nucleic acid sequences, and transgenes are typically integrated into the host genome and expressed in a manner similar to endogenous genes. This difference allows codon optimization in plants to be guided directly by species-specific genomic information.

Plant genomes are highly diverse. At the same time, large scale datasets including pan-genomes, pan-transcriptomes, translatomes and proteomes are rapidly accumulating^20–25^. These datasets now make it feasible to build codon optimization frameworks that can be retrained efficiently using species-specific data. Supervised fine-tuning of large pre-trained biological language models therefore represents a promising strategy for building flexible codon optimization systems.

Here we introduce HalluCodon, a customizable codon optimization framework guided by multimodal biological language models. The framework integrates two complementary modules. The first module, CodonNAT, evaluates the naturalness of coding sequences by measuring how well their codon context matches patterns observed in endogenous genes of the host species. CodonNAT is constructed through joint fine-tuning of the protein language model ESM2 (650M parameters) and the RNA language model mRNA-FM^26, 27^. The model captures sequence features associated with host-specific codon context. In the current implementation, base models were trained using coding sequences from fifteen plant species, including Arabidopsis (*Arabidopsis thaliana*), Canola (*Brassica napus*), Sweet orange (*Citrus sinensis*), Cotton (*Gossypium hirsutum*), Soybean (*Glycine max*), Barley (*Hordeum vulgare*), Medicago (*Medicago truncatula*), Tobacco (*Nicotiana tabacum*), Rice (*Oryza sativa*), Earthmoss (*Physcomitrella patens*), Tomato (*Solanum lycopersicum*), Potato (*Solanum tuberosum*), Wheat (*Triticum aestivum*), Grape (*Vitis vinifera*) and Maize (*Zea mays*). The second module, CodonEXP, predicts the probability that a coding sequence will achieve high protein expression. This model integrates features derived from coding sequences, amino acid sequences and experimental protein abundance data. Prediction accuracy for protein expression reached 82.4% in maize, 82.7% in rice and 84.3% in tobacco, outperforming existing models that rely solely on either protein level or nucleotide level features. These results are consistent with the well-established observation that protein abundance depends not only on physicochemical properties such as folding stability and solubility, but also on nucleotide level features that influence translational efficiency and mRNA stability^28–31^.

Generation of optimized coding sequences can be performed using either genetic algorithms or hallucination-based design, both of which use discriminative models to guide sequence generation. Genetic algorithms have been widely used for designing DNA and protein sequences. In contrast, hallucination-based design has primarily been applied to amino acid sequence generation, for example in *de novo* design of mini protein binders^32–35^. In our benchmarking experiments, hallucination-based design produced higher experimental protein abundance and required substantially less computational time than genetic algorithms. For this reason, hallucination design was adopted as the primary sequence generation strategy in HalluCodon. To facilitate the use of HalluCodon, a web interface (https://codon.oneshot.ac.cn) was developed to allow users to perform codon optimization for the fifteen plant species whose datasets were used to train the base model.

## Results

### An overview of the HalluCodon framework

HalluCodon uses supervised fine-tuning of pre-trained protein (ESM2-650M) and RNA (mRNA-FM) language models to enable highly customizable codon optimization using species-specific datasets provided by users. Features are first extracted from the amino acid (AA) sequences and nucleotide sequences of CDS datasets collected from fifteen plant species using the two language models. The resulting vectors are then integrated through weighted summation to generate merged feature representations of protein and mRNA sequences (**Fig. 1a**). The three sets of vectors (protein, mRNA, and merged) are subsequently used to train a model that predicts codon naturalness (CodonNAT) within a masked language modeling (MLM) framework. Through this training process, the model learns codon context signatures of the host species (**Fig. 1b**). Based on the combined context of codons and amino acids, CodonNAT predicts the codon most likely to occur at each amino acid position. The model then calculates a Naturalness score that quantitatively measures how well a CDS matches the codon context compatibility of a given species. The second module, CodonEXP, predicts codons associated with high protein expression. This module generates a Probability score by linking protein expression data to CDSs in the training dataset (**Fig. 1c**). Protein expression data for the fifteen plant species were obtained from the Protein Abundance Database (PaxDb), where protein abundance values are mapped to the corresponding CDS for each species^36^. For each species, proteins were categorized as High (top 33%) or Low (bottom 33%) expression abundance, and CodonEXP was trained to predict the probability that a CDS corresponds to a protein labeled “High”.

**Fig. 1.**
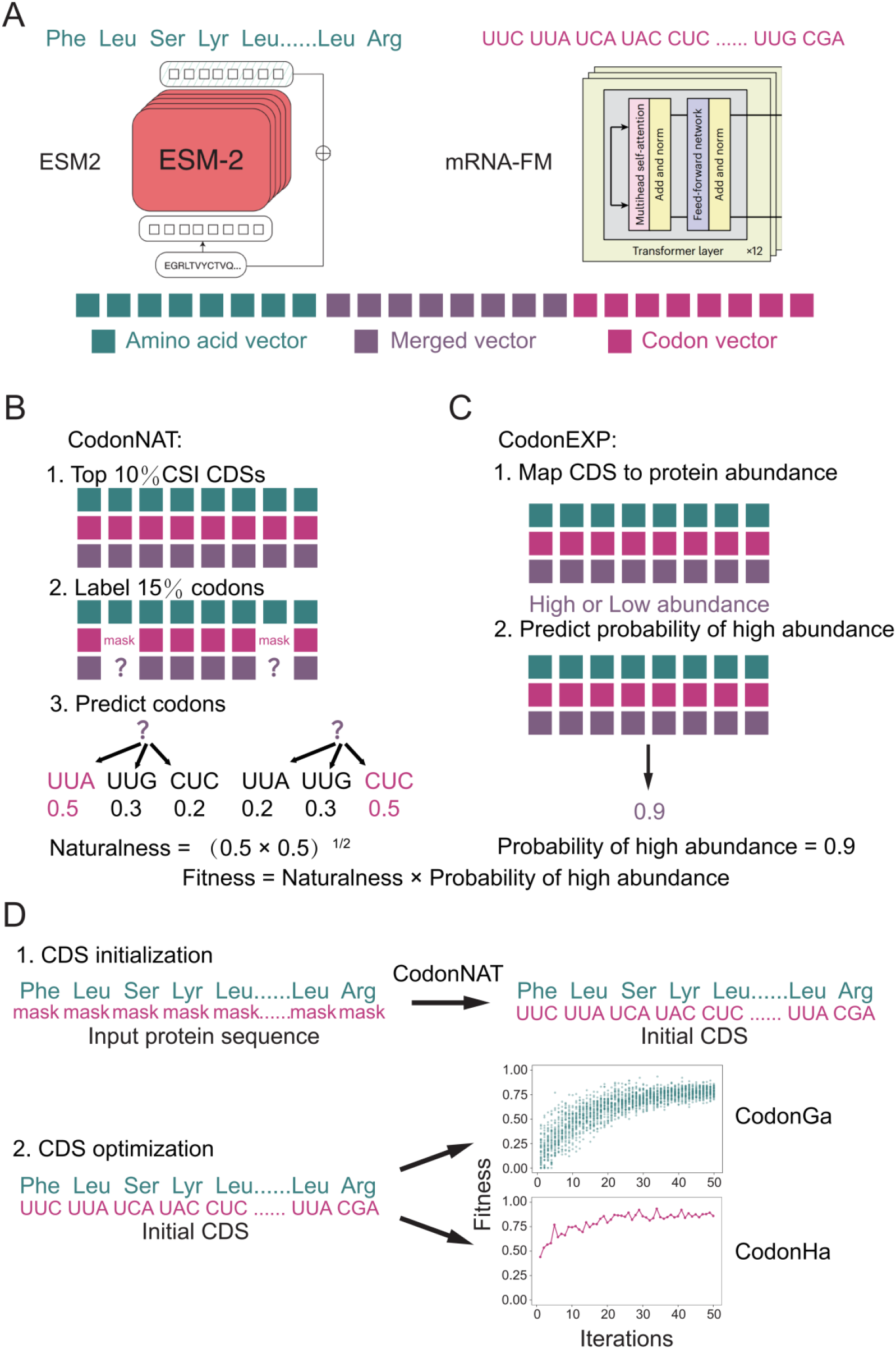
Overview of the HalluCodon framework. **A**. Feature fusion process. Amino acid (AA) vectors and codon vectors are extracted from protein and CDS sequences using the protein language model ESM2 (650M) and the RNA language model mRNA-FM, respectively. The two sets of vectors are combined by weighted summation to generate merged vectors representing integrated protein and CDS features. **B**. Training of the CodonNAT module. CDSs ranked in the top 10% by the codon similarity index (CSI) within the host genome are first selected. A masked language modeling task is then constructed by labeling 15% of codons to learn host-specific codon context features. A Naturalness score is defined as the geometric mean of the predicted probabilities for the labeled codons and is used to quantify the compatibility between the input CDS and the codon context of the host genome. **C**. Training of the CodonEXP module. Each CDS is assigned a binary label indicating High or Low protein expression. CodonEXP predicts the probability of high protein expression given a CDS. A Fitness score is defined as the product of the Naturalness score and the predicted probability of high protein expression and is used to select the optimal CDS. **D**. Two-step generation of optimized CDSs. In the CDS initialization step, the CDS corresponding to the input protein sequence is fully masked at the codon level, after which CodonNAT predicts the most probable synonymous codon for each AA. This step, termed CodonIni, produces an initial CDS for further optimization. In the second step, the CDS is optimized using either a genetic algorithm (CodonGa) or hallucination design (CodonHa). Optimization proceeds through multiple iterations until the Fitness score reaches a plateau, resulting in a final CDS that balances compatibility with host codon usage and a high probability of protein expression.

Generation of a CDS with optimized codon usage must satisfy both codon context compatibility and a high probability of protein expression. This process is implemented in two stages. First, CodonNAT generates an initial CDS from the input protein sequence. The sequence is then iteratively optimized using either a genetic algorithm or hallucination-based design (see **Methods**). During each iteration, a Fitness score is calculated as the product of the Naturalness score from CodonNAT and the Probability score predicted by CodonEXP. This Fitness score guides the optimization process and determines the final CDS (**Fig. 1d**).

### CodonNAT infers the compatibility of codon context with the host genome

The CodonNAT model learns codon context signatures from host genomes and evaluates the compatibility of a CDS within a given species. Model training and evaluation followed the strategy used in CodonTransformer^14^. First, non-redundant CDSs were ranked using the codon similarity index (CSI), and the top 10% were selected from the reference genome annotations of fifteen plant species (ranging from 3539 genes in sweet orange to 13,438 genes in wheat; **Supplementary table 1**). From this dataset, 90% of the CDSs were used for model training and validation. The remaining 10% were reserved for independent testing. To avoid sequence redundancy, test sequences showing more than 90% similarity to any training sequence were removed using CD-HIT. Model accuracy was calculated as the frequency of correctly predicted original codons at masked positions. Because the RNA language model mRNA-FM accepts a maximum input length of 3,066 nucleotides, only the first 1,022 codons of each CDS were used during CodonNAT training. Cross-validation showed prediction accuracies of 77.0% in maize, 78.9% in rice, and 63.9% in tobacco (**Fig. 2a**). Across species, the average prediction accuracy of CodonNAT was 66.5%, substantially higher than the 56.6% achieved by the traditional BFC (background frequency choice) method, which always selects the most frequent codon for a given amino acid within a species. CodonNAT also outperformed BFC for nearly all twenty amino acids across species (**Supplementary Fig. 1**).

**Table 1.** Performance comparison (mean ± s.d.) of CodonEXP and other language models in predicting protein expression across maize, rice, and tobacco.

| Species | Model | AUC | ACC | Recall | Precision | F1 | MCC |
| --- | --- | --- | --- | --- | --- | --- | --- |
| Maize | CodonEXP | 0.88±0.017 | **0.82±0.006** | **0.85±0.025** | 0.81±0.016 | **0.83±0.007** | **0.65±0.013** |
| | ESM2 | **0.89±0.006** | 0.82±0.012 | 0.83±0.023 | **0.81±0.028** | 0.82±0.008 | 0.64±0.022 |
| | mRNA-FM | 0.87±0.014 | 0.81±0.007 | 0.84±0.014 | 0.79±0.017 | 0.81±0.003 | 0.62±0.012 |
| | CaLM | 0.85±0.006 | 0.78±0.002 | 0.80±0.044 | 0.77±0.022 | 0.78±0.010 | 0.55±0.005 |
| | PlantRNA-FM | 0.76±0.068 | 0.68±0.066 | 0.76±0.093 | 0.67±0.070 | 0.71±0.051 | 0.38±0.120 |
| | RNA-FM | 0.59±0.105 | 0.55±0.074 | 0.87±0.178 | 0.55±0.066 | 0.66±0.020 | 0.11±0.147 |
| Rice | CodonEXP | **0.90±0.017** | **0.83±0.002** | **0.88±0.036** | 0.79±0.020 | **0.83±0.005** | **0.66±0.005** |
| | ESM2 | 0.89±0.006 | 0.82±0.010 | 0.85±0.025 | **0.80±0.023** | 0.82±0.008 | 0.65±0.018 |
| | mRNA-FM | 0.88±0.016 | 0.81±0.010 | 0.88±0.034 | 0.77±0.026 | 0.82±0.005 | 0.62±0.014 |
| | CaLM | 0.86±0.006 | 0.78±0.007 | 0.82±0.027 | 0.75±0.019 | 0.78±0.006 | 0.56±0.012 |
| | PlantRNA-FM | 0.71±0.129 | 0.65±0.102 | 0.83±0.094 | 0.62±0.089 | 0.70±0.042 | 0.30±0.198 |
| | RNA-FM | 0.55±0.124 | 0.53±0.083 | 0.96±0.088 | 0.52±0.065 | 0.67±0.023 | 0.07±0.166 |
| Tobacco | CodonEXP | **0.91±0.012** | **0.84±0.013** | **0.86±0.021** | 0.81±0.018 | **0.83±0.014** | **0.69±0.027** |
| | ESM2 | 0.89±0.007 | 0.83±0.008 | 0.82±0.009 | **0.81±0.010** | 0.82±0.008 | 0.66±0.015 |
| | mRNA-FM | 0.89±0.011 | 0.81±0.009 | 0.86±0.035 | 0.76±0.013 | 0.81±0.012 | 0.64±0.020 |
| | CaLM | 0.85±0.010 | 0.77±0.016 | 0.81±0.022 | 0.72±0.033 | 0.76±0.008 | 0.54±0.027 |
| | PlantRNA-FM | 0.77±0.032 | 0.71±0.021 | 0.71±0.076 | 0.68±0.041 | 0.69±0.027 | 0.42±0.039 |
| | RNA-FM | 0.77±0.020 | 0.70±0.030 | 0.69±0.053 | 0.67±0.052 | 0.68±0.013 | 0.40±0.049 |

**Fig. 2.**
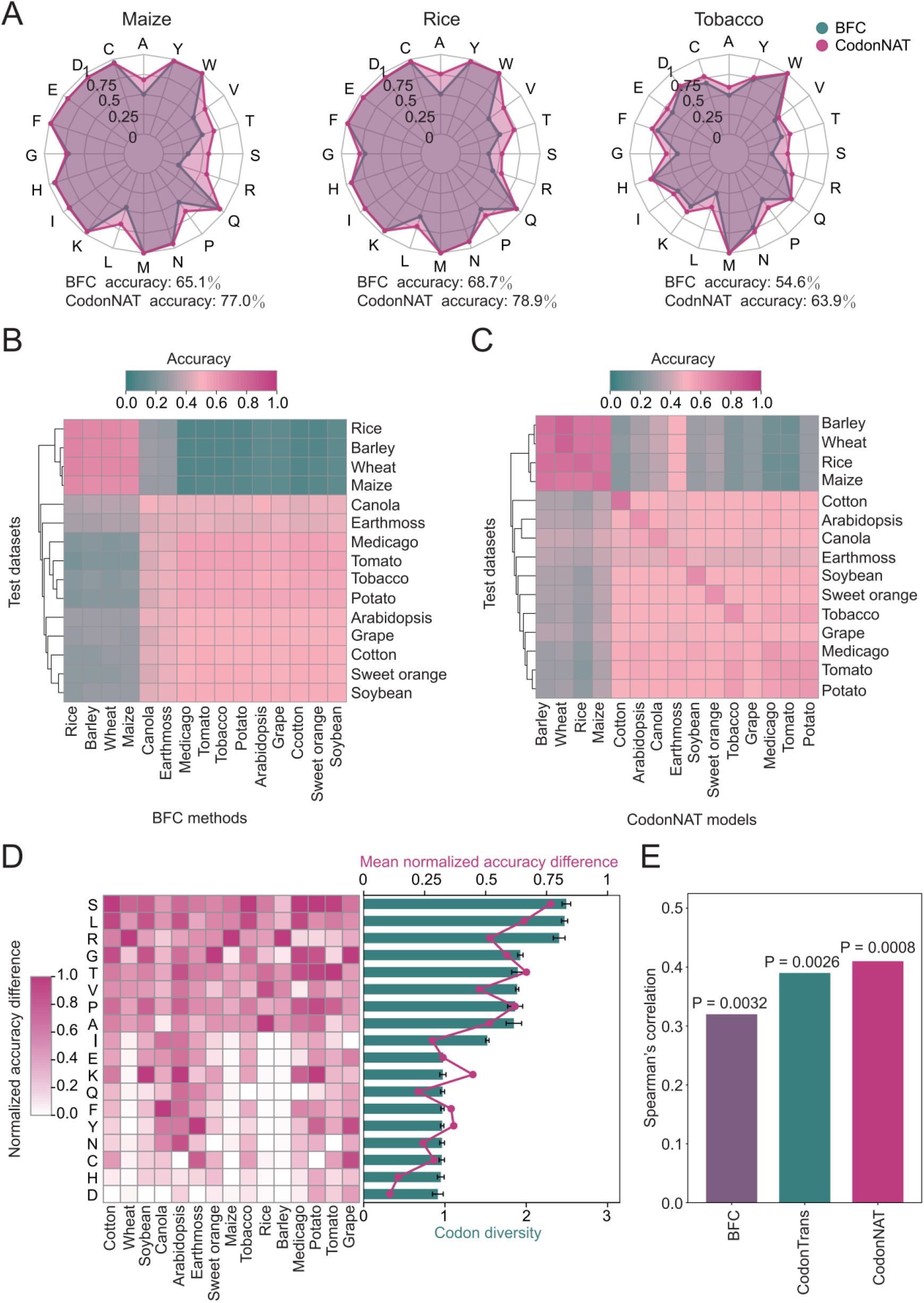
CodonNAT learns codon context signatures of the host genome. **A**. Prediction accuracy of synonymous codons for the 20 amino acids (AAs) in maize, rice, and tobacco. **b, c**. Clustering of codon usage patterns across the fifteen plant species based on prediction results from the BFC (b) and CodonNAT (c) methods. **d**. Relationship between codon diversity and the mean normalized accuracy difference between CodonNAT and BFC predictions. Codon diversity is defined as the Shannon entropy of the synonymous codon usage frequency distribution. The mean normalized accuracy difference is a value between 0 and 1, calculated by applying Min-Max normalization to the accuracy differences of all AAs within each species, followed by averaging the normalized values for each AA across species. **e**. Comparison of BFC, CodonTransformer, and CodonNAT models based on the correlation between their predicted scores and experimentally measured fitness values for 62 synonymous mutations in the *Escherichia coli ccdA* gene. Correlations were evaluated using two-tailed Spearman’s rank correlation tests.

To further examine differences in codon usage among species, we evaluated codon prediction accuracy across all datasets using both the BFC method and the CodonNAT model. Clustering of the fifteen species based on prediction accuracy (see **Methods**) grouped monocots (Rice, Barley, Wheat, Maize) together, while dicots formed a separate cluster under both approaches (**Fig. 2a** and **2b**). For example, the BFC model trained on rice achieved prediction accuracies of 69.1%, 69.1%, and 66.9% for Barley, Wheat, and Maize, respectively, but only 19.7% for Tomato. Similarly, the CodonNAT model trained on rice achieved accuracies of 75.3%, 74.9%, and 72.6% for Barley, Wheat, and Maize, but only 22.9% for Tomato. These results suggest that codon usage patterns reflect phylogenetic relationships among plant species. Interestingly, a CodonNAT model trained on the basal terrestrial plant *Physcomitrella patens* (Earthmoss) achieved prediction accuracies of 50.8%, 51.8%, 51.7%, and 50.6% for rice, barley, wheat, and maize, respectively, whereas the corresponding BFC method achieved only 25.9%, 26.9%, 27.2%, and 26.7%. This result indicates that CodonNAT captures evolutionarily conserved codon usage features beyond simple codon frequency. We next examined how prediction performance varies among amino acids. For each species, we calculated the difference in synonymous codon prediction accuracy between CodonNAT and BFC for each amino acid. To remove species-specific effects, these differences were normalized using Min–Max normalization within each species. We then calculated the mean normalized accuracy difference for each amino acid across species. Amino acids with higher codon diversity, measured using Shannon entropy, tended to show larger accuracy improvements for CodonNAT (see **Methods**). This observation indicates that CodonNAT learns sequence information beyond simple codon usage frequencies and performs particularly well at positions with high codon diversity (**Fig. 2d**).

Following the benchmarking strategy used in CodonTransformer, we performed an additional analysis using the *ccdA* benchmark gene in *Escherichia coli* (see **Method**). The *ccdA* gene encodes the antitoxin component of the *ccdAB* toxin– antitoxin system in *E. coli*, and experimental fitness values are available for 62 synonymous mutations^37^. This dataset therefore provides a useful benchmark for assessing the relationship between codon usage and cellular fitness. After training CodonNAT on CDSs corresponding to the top 10% CSI genes in *E. coli* (K12 strain), the model was able to predict the fitness effects of the 62 synonymous mutations using a zero-shot learning approach. We compared predictions generated by BFC, CodonTransformer, and CodonNAT. Spearman correlation coefficients between predicted and measured fitness values were 0.32, 0.39, and 0.41, respectively (**Fig. 2e**). These results indicate that CodonNAT achieved the highest correlation among the three models, supporting its ability to capture biologically meaningful codon usage patterns.

### CodonEXP infers the probability of high protein expression

Protein abundance varies substantially among endogenous genes within a host species. Predicting protein expression levels directly from CDS features is therefore an important component of effective codon optimization. CodonEXP was designed to estimate the probability that a CDS will produce high levels of protein expression in a given host species. To evaluate CodonEXP, we used a dataset of artificially constructed codon-randomized monomeric red fluorescent proteins (mRFPs), which includes 1,459 variants with different CDSs. In this dataset, synonymous codons were randomized across the entire CDS without altering the amino acid sequence, and protein abundance of each mRFP variant was measured after expression in *E. coli*^38^. The 1,459 variants were divided into training and testing sets at a ratio of 4:1. Because all variants share the same amino acid sequence, protein-level features remain constant. Therefore, the weights of the ESM2 component in CodonEXP were fixed, and the original ESM2 model was used to extract protein sequence representations. Five-fold cross-validation (CV) was performed on the training set, generating five models that predicted the probability of high mRFP expression. We then calculated the Spearman correlation between predicted and observed expression levels on the test set. To benchmark CodonEXP, we compared its performance with four recently developed RNA language models: RNA-FM, PlantRNA-FM, CaLM, and mRNA-FM^27, 39, 40^. These models represent state-of-the-art approaches for capturing RNA features associated with protein abundance, including mRNA stability, translation efficiency, and RNA secondary structure. Using the same training–testing splits, these models were trained and evaluated with pooling layers and regression heads identical to those used in CodonEXP. Evaluation on the testing set showed that mRNA-FM achieved the best performance among the RNA language models, with an average Spearman correlation of 0.84 (**Fig. 3a**). PlantRNA-FM showed the lowest performance (*r* = 0.25), likely because it was pre-trained exclusively on plant RNA sequences and therefore lacks compatibility with non-plant systems such as *E. coli* used in this evaluation. CodonEXP achieved a Spearman correlation of 0.85, slightly outperforming mRNA-FM. The final predicted expression values for the 292 CDSs in the testing set were obtained by averaging predictions from the five models generated during cross-validation. The coefficient of determination (*r*^2^) between these averaged predictions and the measured mRFP abundances reached 0.75 (**Fig. 3b**).

**Fig. 3.**
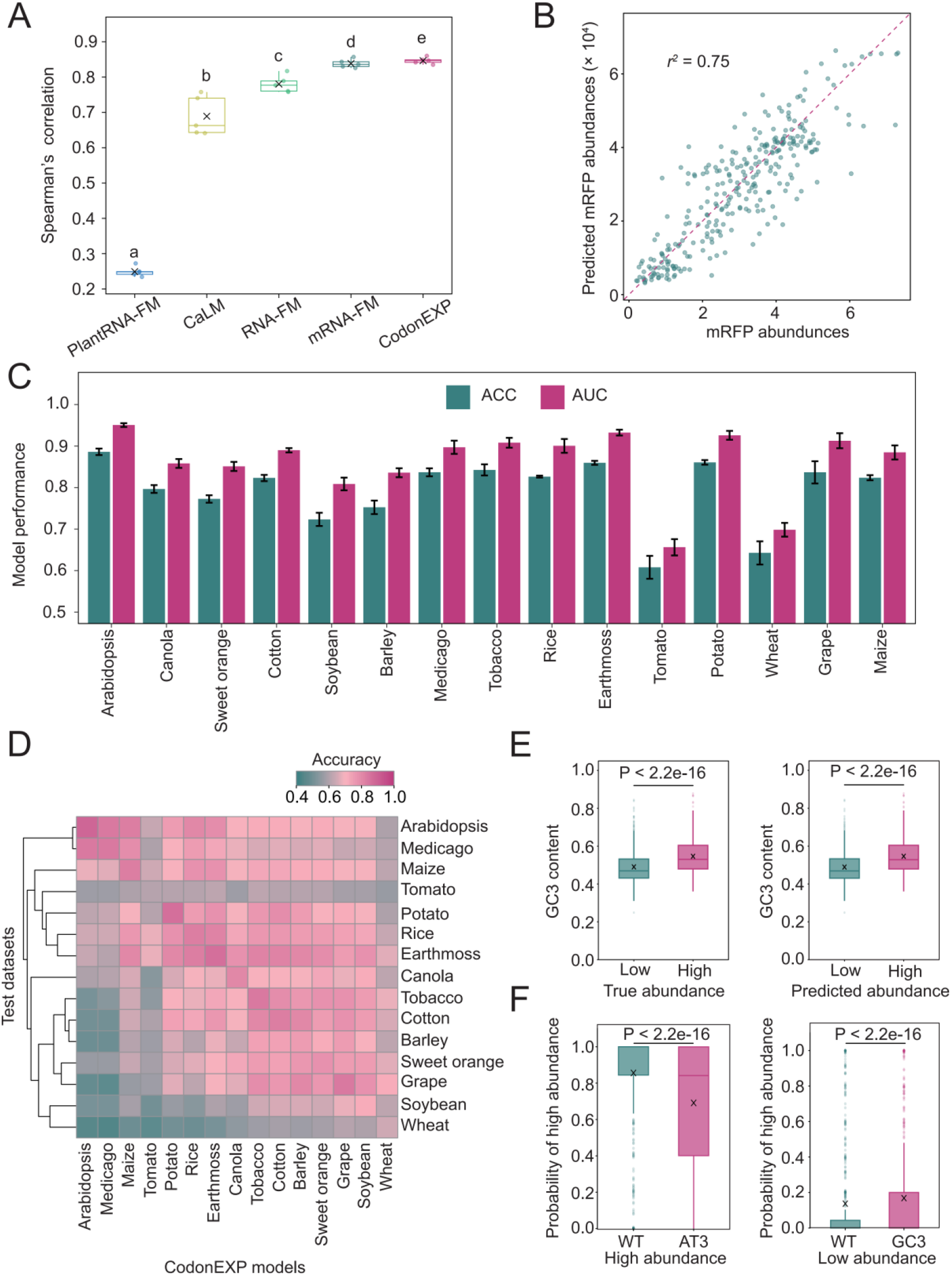
CodonEXP infers the probability of high protein expression. **A**. CodonEXP outperforms other models when evaluated using experimentally measured mRFP expression data in *E. coli*. Letter annotations (a–e) indicate statistically significant differences between groups (*p* < 0.05, Friedman test followed by Holm-adjusted Conover post hoc multiple comparisons). **b**. Coefficient of determination (*r*^2^ = 0.75) between CodonEXP predictions and experimentally measured relative abundance of mRFP protein. **c**. Evaluation of CodonEXP performance across the fifteen plant species using ACC and AUC metrics. Error bars represent standard deviations from five-fold cross-validation. **d**. Cross-species prediction performance of CodonEXP. **e**. Comparison of GC3 content between high- and low-abundance proteins in Earthmoss. Statistical significance was assessed using the Mann–Whitney test, with exact p-values indicated above the boxes. Each boxplot shows group mean values (marked with an X) and the corresponding GC3 distribution. **f**. Comparison of predicted probabilities of high abundance by CodonEXP between original CDSs and synonymous codon variants with altered GC3 content in Earthmoss. Statistical significance was assessed by the paired Wilcoxon signed-rank test, with exact p-values indicated above the boxes. Each boxplot displays group mean values (marked with an X) and corresponding predicted probabilities of high abundance.

Predicting protein expression through direct regression often suffers from substantial noise in public datasets. We therefore followed the strategy proposed by Liu et al. and formulated the problem as a classification task that predicts the probability of high protein expression^28^. For the fifteen plant species with available protein abundance data in PaxDb, CDSs were labeled as High (top 33%) or Low (bottom 33%) based on their protein abundance levels (see **Methods**). After clustering protein sequences using CD-HIT with a 90% identity threshold, the remaining CDSs were divided into training and testing sets at a ratio of 4:1. Unlike the mRFP dataset, protein sequences in this dataset vary substantially. Therefore, the weights of the ESM2 component in CodonEXP were not fixed during training. This allowed the model to learn informative sequence representations from both CDS and amino acid sequences for improved prediction of protein abundance labels. Five-fold cross-validation was performed on the training set for each species, generating five models that predicted the probability of high expression. Model performance was evaluated on the testing sets using several metrics, including area under the receiver operating characteristic curve (AUC), accuracy (ACC), recall, precision, F1 score (F1), and Matthews correlation coefficient (MCC). Across the fifteen plant species, CodonEXP achieved average ACC and AUC values of 79.3% and 86.1%, respectively (**Fig. 3c, Supplementary Fig. 2 and 3**). We further compared CodonEXP with the four RNA language models described above using datasets from Maize, Rice, and Tobacco. For a fair comparison, all models were trained using the same pooling layers and classification heads as CodonEXP. CodonEXP consistently outperformed the other models across all evaluation metrics and datasets, indicating that it successfully integrates information from both CDS and amino acid sequences (**Table 1**). Notably, CodonEXP also surpassed the ESM2 model alone in terms of ACC, recall, F1, and MCC across all three species, suggesting that additional information encoded in CDS sequences contributes to improved prediction of protein expression. This result is consistent with previous observations that protein abundance depends not only on protein physicochemical properties but also on nucleotide-level features affecting translation efficiency and mRNA stability^28–31^.

To examine the cross-species predictive capability of CodonEXP, we used the five models trained through cross-validation in each species to predict the probability of high protein expression across all fifteen species. For each evaluation, sequences with high similarity to training data were removed from the testing sets (see **Methods**). Prediction performance varied among species, reflecting differences in cellular environments and codon usage patterns (**Fig. 3d**). Nevertheless, CodonEXP maintained reasonable predictive accuracy even when applied to species not included in the training data, demonstrating strong generalization capability. For example, the CodonEXP model trained on the basal terrestrial plant Earthmoss achieved an average prediction accuracy of 70.6% across test sets from the remaining fourteen species. This result indicates that CodonEXP captures conserved sequence features associated with high protein expression across species. Furthermore, both CodonEXP predictions and experimentally derived labels showed significantly higher GC3 content in CDSs corresponding to highly expressed proteins compared with low-abundance proteins in Earthmoss (**Fig. 3e**). This observation is consistent with previous reports that plant genes with high translation efficiency often show enrichment of G or C at the third codon position (GC3), a pattern successfully captured by CodonEXP. Next, for the high-expression CDSs in the Earthmoss test data, we replaced all codons with synonymous codons ending with A or T (AT3) whenever possible. As a result, CodonEXP predicted a significant decrease in the Probability score of high protein expression. Conversely, for the low-expression CDSs, we replaced all codons with synonymous codons ending with G or C (GC3) whenever possible, which led to a significant increase in the predicted Probability score of high protein expression by CodonEXP (**Fig. 3f**). We observed a similar pattern in other plant species as well, indicating that CodonEXP captures the important role of GC3 in enhancing protein expression (**Supplemental Fig. 4**).

### CDS generation with genetic algorithms and hallucination design

Both genetic algorithms and hallucination design can use discriminative models to guide generative sequence design. Genetic algorithms have been widely applied to the design of DNA and amino acid sequences^32, 33^. Hallucination-based design, in contrast, has mainly been used to generate protein sequences, for example in the *de novo* design of proteins predicted to fold into specific backbone structures or in the design of mini-protein binders targeting a protein of interest^34, 35^. In HalluCodon, the iterative optimization of CDS sequences is guided by a Fitness score. This optimization can be implemented using either a genetic algorithm, referred to as CodonGa, or a hallucination-based algorithm, referred to as CodonHa. In both cases, the Fitness score is defined as the product of the Naturalness score predicted by CodonNAT and the Probability score of high protein expression predicted by CodonEXP.

To examine sequence patterns generated during iterative optimization, we applied both algorithms to optimize the CDS of the DsRed2 gene in Tobacco, which encodes a red fluorescent protein widely used as a marker in cellular imaging experiments^41^. When CodonGa was used for CDS generation, the optimal DsRed2 sequence was obtained after the 53^rd^ iteration, reaching a maximum Fitness score of 0.6598 (**Fig. 4a**). Using CodonHa, the optimal sequence was generated after only 13 iterations, with a maximum Fitness score of 0.6237 (**Fig. 4b**). During the optimization process, the Naturalness score remained largely stable for both algorithms, whereas the Probability score for high protein expression rapidly approached 1.0. This occurred after approximately 21 iterations with CodonGa and after only four iterations with CodonHa (**Supplementary Fig. 5**). These results suggest that the optimization procedure primarily improves predicted expression probability while maintaining compatibility with host codon context. CodonHa also showed substantially higher computational efficiency. On an RTX3090 GPU, CodonHa completed codon optimization in 138.22 seconds, which is 46.8 times faster than CodonGa, which required 6465.74 seconds (**Fig. 4a** and **4b**). In addition, previous studies have shown that increased usage of codons with high GC3 content can enhance both mRNA abundance and protein production^42^. Consistent with this observation, the GC3 content of optimized CDS sequences increased markedly during CodonHa optimization, whereas CodonGa showed a slower and more gradual increase during the optimization process (**Fig. 4c** and **4d**). Together, these results indicate that the hallucination-based CodonHa approach provides a more computationally efficient strategy for CDS optimization than the genetic algorithm–based CodonGa method.

**Fig. 4.**
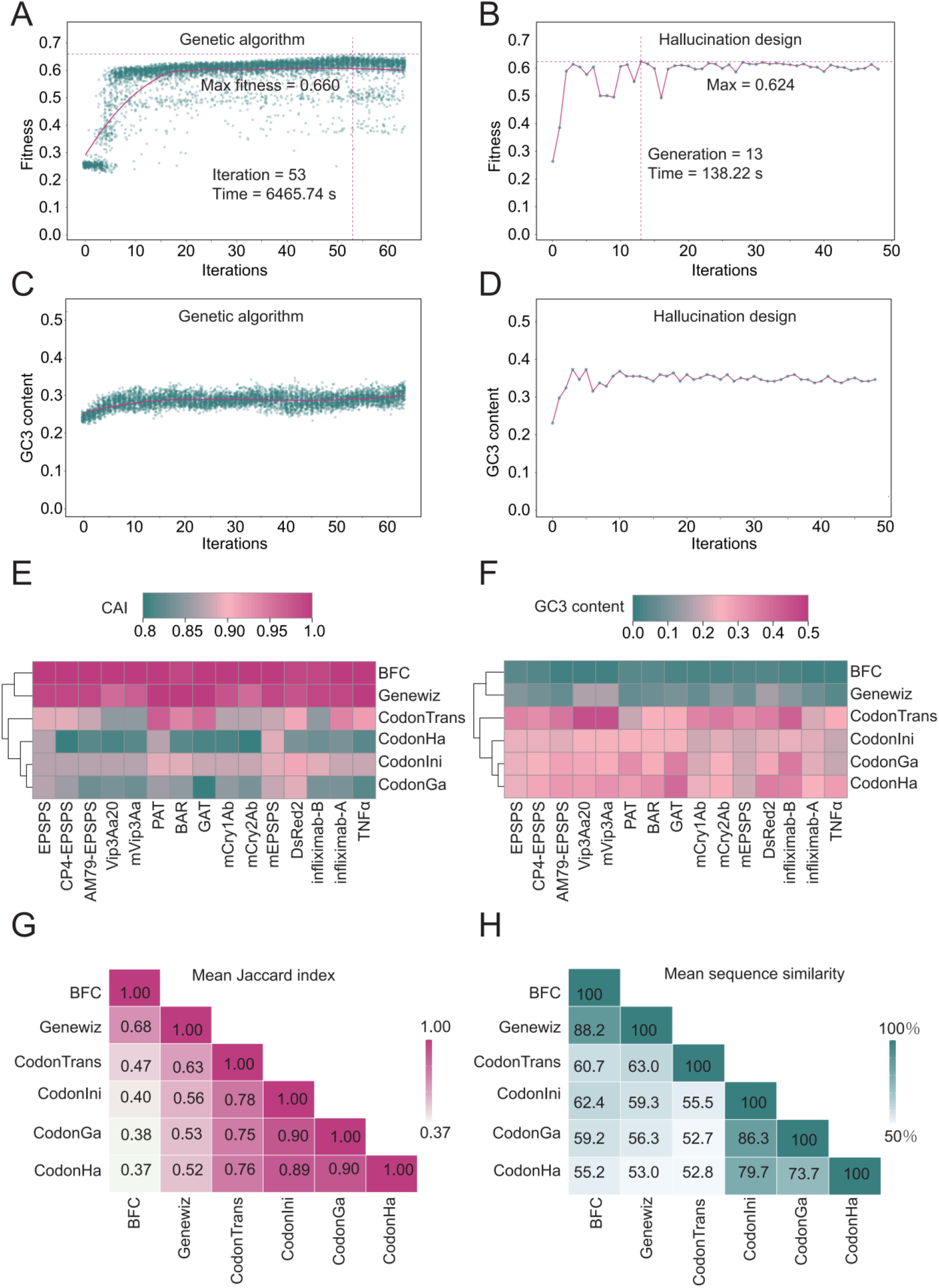
Comparison of different CDS generation methods. **a, b**. Fitness scores increase and reach plateaus during iterative optimization of the DsRed2 CDS using the CodonGa (a) and CodonHa (b) algorithms. **c, d**. GC3 content increases and reaches plateaus during iterative optimization of the DsRed2 CDS using the CodonGa (c) and CodonHa (d) algorithms. Green dots represent the Fitness score or GC3 content of each optimized CDS throughout the iterative process. The red line shows the overall trend of Fitness score or GC3 content, fitted using locally weighted scatterplot smoothing (LOESS). **e, f**. Clustered patterns of CAI (e) and GC3 content (f) calculated from the optimized CDSs of the fifteen benchmark proteins generated by the six methods. CodonTrans denotes the CodonTransformer method. **g, h**. Mean Jaccard index (g) and mean sequence similarity (h) among optimized CDSs of the fifteen benchmark proteins generated by different methods.

### Comparison of multiple codon optimizers using benchmark proteins

To compare the performance of HalluCodon with existing codon optimization tools, we selected fifteen proteins as benchmarks (**Supplementary dataset 1**). These proteins are widely used in molecular experiments (for example DsRed2), transgenic crop development (EPSPS, mCry2Ab, Vip3Aa20, among others), or have potential applications in plant molecular farming (infliximab-A and infliximab-B). Three codon optimization methods—BFC, Genewiz, and CodonTransformer—were compared with the two-step optimization procedure implemented in HalluCodon for Tobacco. In the first step, the CodonNAT module predicts codons for a given protein sequence by selecting the most probable codon for each amino acid. The resulting sequence represents an initial CDS and this step is referred to as CodonIni. In the second step, the initial CDS is further optimized by iteratively maximizing the Fitness score using either the CodonGa (genetic algorithm) or CodonHa (hallucination-based) methods.

We first evaluated the codon adaptation index (CAI) of CDS sequences generated for the fifteen benchmark proteins by each method (**Fig. 4e**). CAI measures the relative usage frequency of preferred codons in a species, with values ranging from 0 to 1, representing the lowest to highest levels of codon adaptation. The BFC method generated CDS sequences with a CAI of 1.0 for all fifteen proteins, consistent with the fact that BFC selects the most frequent codon for each amino acid in the host species. Genewiz produced CDS sequences with an average CAI of 0.98, indicating that its algorithm also relies largely on codon usage frequency. In contrast, CodonTransformer, CodonIni, CodonGa, and CodonHa produced CDS sequences with average CAI values of 0.90, 0.88, 0.84, and 0.83, respectively. Although language model–based approaches generated slightly lower CAI values, the resulting CDS sequences still maintained relatively high codon adaptation.

We next examined the GC3 content of codons optimized by the six methods. Among the top 10% CSI CDS sequences in tobacco, the natural GC3 frequency is 0.31. However, the BFC method generated optimized CDS sequences with an average GC3 content of only 0.03, far below the observed genomic frequency. CodonIni increased the average GC3 content to 0.23 compared with BFC, although this value remained lower than the natural GC3 frequency. CodonGa further increased GC3 content to 0.27. CodonHa achieved an average GC3 content of 0.30, which is closest to the natural GC3 frequency (0.31) observed in tobacco among the six methods. CodonTransformer produced the highest GC3 content (0.34) (**Fig. 4f**).

We also compared the codon usage similarity of CDSs encoding the same fifteen benchmark proteins generated by the six optimization methods. Codon usage similarity was calculated using the Jaccard index, defined as the ratio of the intersection to the union of codon sets between two CDS sequences^43^. Values closer to 0 indicate lower codon usage similarity, whereas values closer to 1 indicate higher similarity. Among the tested methods, CodonTransformer showed the highest average similarity (0.76) with HalluCodon-generated sequences. Genewiz and BFC ranked second (0.63) and third (0.47), respectively (**Fig. 4g**). This observation likely reflects the fact that both CodonTransformer and HalluCodon are based on biological language models, whereas Genewiz and BFC primarily rely on codon frequency statistics. Notably, although both HalluCodon and CodonTransformer were trained using the same CDS dataset from Tobacco, the sequence similarity between optimized CDS sequences generated by the two methods was only about 53.7%. In addition, CodonGa and CodonHa shared mean sequence similarities of 86.3% and 79.7%, respectively, with the initial CDS generated by CodonIni (**Fig. 4h**). These results suggest that CodonHa explores a broader CDS sequence space during optimization and may therefore achieve better compatibility with host codon usage patterns than CodonGa.

### Experimental validation of optimized proteins

To experimentally validate the performance of HalluCodon, we selected five proteins: DsRed2, mCry2Ab, glyphosate acetyltransferase (GAT), infliximab-A, and infliximab-B and tested their expression in Tobacco leaves using a transient expression system (**Supplementary dataset**). For DsRed2, CDS sequences optimized by the six methods were synthesized and transiently expressed in Tobacco leaves, and fluorescence intensity was measured as a proxy for protein expression (**Fig. 5a**). The DsRed2 sequence optimized by CodonHa produced the strongest mRFP fluorescence signal. The red fluorescence intensity was 1.57 times higher than that obtained with CodonTransformer, 4.32 times higher than with Genewiz, and 13.58 times higher than with BFC. These results were consistent with Western blot analysis of DsRed2 protein expression (**Fig. 5b** and **5c, Supplementary Fig. 6a**). One possible explanation for the higher protein yield is the increased GC3 content in sequences optimized by CodonTransformer and CodonHa, both of which reached a GC3 value of 0.36. However, despite having similar GC3 content, CodonHa produced higher expression levels, suggesting that the algorithm may capture additional features of codon context beyond GC3 enrichment that contribute to increased protein production. A similar pattern was observed for GAT. The CDS optimized by CodonHa showed the strongest protein band in Western blot analysis, while CodonTransformer ranked second (**Fig. 5c, Supplementary Fig. 6b**). Together, these results support the superior performance of CodonHa, likely due in part to its enrichment of GC3 codons, which has been associated with improved mRNA stability and increased protein production^42^. Based on these observations, we hypothesized that further increasing GC3 content in addition to CodonHa optimization might enhance protein production. To test this hypothesis, we artificially increased the GC3 content of CDSs for DsRed2, GAT, mCry2Ab, infliximab-A, and infliximab-B to 0.8 using the so-called “GC3 method”, implemented through the online tool GC3-recoder^42^. This modification increased protein abundance to levels comparable to those obtained with CodonHa-optimized sequences (**Fig. 5d–g, Supplementary Fig. 6c–e**). Notably, large proteins such as mCry2Ab and infliximab-B failed to express when optimized solely with the original CodonHa method (**Supplementary Fig. 6e and 6f**). In contrast, when GC3 rewarding was incorporated into the CodonHa optimization process (Ha-GC3; **see Method**), both proteins were successfully expressed (**Fig. 5f and 5g**). These results suggest that increased GC3 content may be particularly beneficial for expressing large proteins, possibly because longer mRNA transcripts require higher GC3 levels to maintain sufficient mRNA stability. However, although elevated GC3 content can improve protein expression, excessively high GC3 levels introduced through random codon substitution across entire CDS also increase overall GC content. This increase can complicate gene synthesis and cloning and may substantially increase the number of CG and CHG methylation sites, by approximately two-to three-fold when a fixed GC3 value of 0.8 is applied (**Supplementary Fig. 7**)^44, 45^. Higher numbers of methylation sites may increase the likelihood of DNA methylation within coding regions, which can lead to transcriptional repression^46^. To avoid arbitrary increases in GC3 content, the GC3-rewarding strategy was integrated directly into the CodonHa optimization algorithm. In this approach, GC3 codons are introduced selectively at appropriate positions during CDS optimization. This allows the algorithm to maintain relatively moderate but optimized GC3 levels while still achieving efficient protein expression, as illustrated in Fig. 5h.

**Fig. 5.**
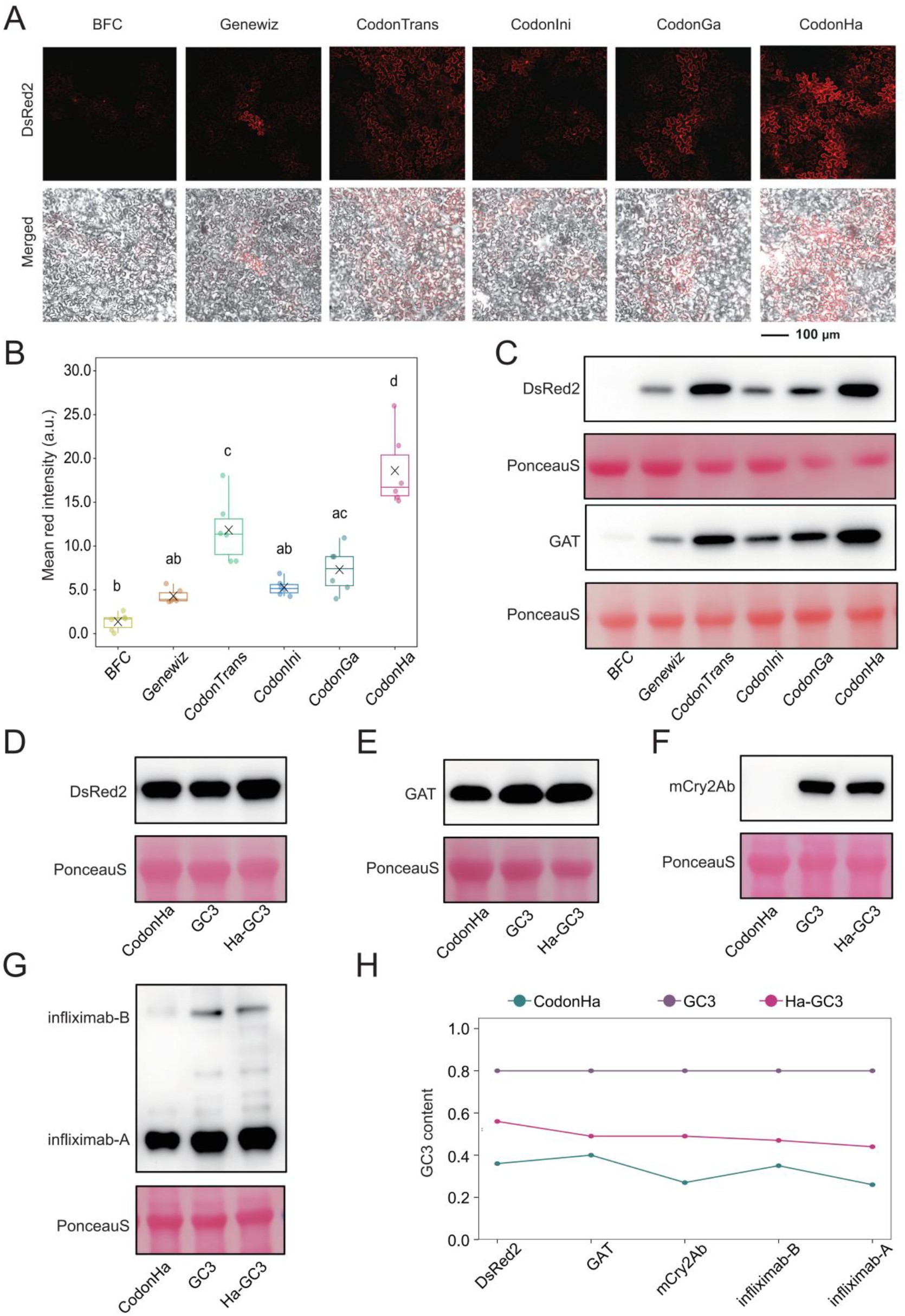
Experimental validation of five benchmark proteins with optimized CDSs in tobacco. **A**. Observation of mRFP fluorescence in tobacco leaves transiently expressing DsRed2 CDSs optimized by the six methods. **b**. Quantitative analysis of mRFP fluorescence intensity based on the images in (a). Six random fields of view were selected for each sample, and red fluorescence intensity was measured using ImageJ. One-way ANOVA followed by Tukey’s HSD test was used for multiple comparisons. Different letter annotations indicate statistically significant differences between groups (*p* < 0.05). **c**. Western blot analysis of DsRed2 and GAT protein abundance using CDSs optimized by the six methods. **d – f**. Western blot analysis of benchmark proteins optimized using the CodonHa, GC3, and Ha-GC3 methods. **h**. Comparison of GC3 content in CDSs optimized by the three methods.

## Discussion

HalluCodon was developed to support diverse applications in which plant cells are used to express heterologous proteins. Such applications include transgenic crop development, plant synthetic biology, and molecular farming of pharmaceutical or industrial proteins. In these contexts, optimized coding sequences can improve protein yield and stability of transgene expression. In addition to optimizing exogenous genes, HalluCodon may also be applied to endogenous genes with agronomic importance. In such cases, modest increases in protein expression could allow fine-tuning of desirable traits without altering regulatory DNA sequences or amino acid composition. This possibility may be particularly valuable in breeding programs, where small quantitative changes in protein abundance can influence trait performance.

A key feature of HalluCodon is its flexible and customizable fine-tuning framework. Users can retrain species-specific models using publicly available datasets or their own experimental data, allowing the method to be adapted to a wide range of research and breeding applications. This flexibility is particularly advantageous for codon optimization in plants. Unlike animal systems, where optimization must account for immune responses to exogenous RNA sequences, plants generally do not mount immune responses against foreign nucleic acids^17^. As a result, codon optimization strategies guided by species-specific genomic data may be especially effective in plant systems. The rapid expansion of plant omics resources, including pan-genomic, pan-transcriptomic, translatomic, and proteomic datasets, further increases the potential of such approaches^20–25^. HalluCodon combines pre-trained protein and RNA language models with experimental labels related to protein expression, allowing the model to learn sequence features associated with efficient translation and protein production. As additional datasets become available, the predictive performance of these models is expected to improve further. In addition, the hallucination-based optimization strategy uses gradient information from discriminative models to guide iterative sequence design toward improved expression. With sufficiently rich training data, these discriminative models can capture multiple determinants of gene expression, including protein abundance, mRNA stability, and translation efficiency. In principle, this framework could support multi-objective optimization in which several expression-related properties are optimized simultaneously.

Our results also highlight the ability of HalluCodon to identify sequence features associated with codon context in plant genomes. Previous studies have reported that GC content at the third codon position (GC3) correlates positively with protein expression levels in plants, likely through effects on mRNA stability and translation efficiency^42^. During CDS optimization with HalluCodon, GC3 content increased in the optimized sequences, indicating that the model captures this characteristic feature of plant codon usage. This observation prompted us to examine whether further enrichment of GC3 codons could enhance protein expression. Experimental validation suggested that GC3 enrichment can indeed improve expression, particularly for large proteins. One possible explanation is that increased GC3 content stabilizes longer mRNA transcripts, thereby supporting higher protein production. However, excessive GC3 enrichment can also increase overall GC content within coding regions. This increase may complicate gene synthesis and cloning and can substantially increase the number of CG and CHG methylation sites^44–46^. Higher densities of methylation sites raise the risk of DNA methylation within coding regions, which may lead to transcriptional repression. Importantly, HalluCodon avoids this problem by balancing GC3 enrichment during optimization rather than maximizing it indiscriminately. In our experiments, the algorithm maintained GC3 frequencies within a moderate range while still achieving high protein expression levels. This behavior suggests that the model captures additional sequence features that contribute to efficient expression beyond GC3 content alone. Such balanced optimization may therefore be advantageous in practical applications, as it improves protein expression while minimizing potential negative effects associated with strongly biased codon composition.

## Methods

### Compilation of training and testing datasets

To train CodonNAT, CDSs were collected from the reference genomes of 15 plant species: Arabidopsis (*Arabidopsis thaliana*), Canola (*Brassica napus*), Sweet orange (*Citrus sinensis*), Cotton (*Gossypium hirsutum*), Soybean (*Glycine max*), Barley (*Hordeum vulgare*), Medicago (*Medicago truncatula*), Tobacco (*Nicotiana tabacum*), Rice (*Oryza sativa*), Earthmoss (*Physcomitrella patens*), Tomato (*Solanum lycopersicum*), Potato (*Solanum tuberosum*), Wheat (*Triticum aestivum*), Grape (*Vitis vinifera*), and Maize (*Zea mays*). CDSs were filtered to retain only those whose lengths were multiples of three and that lacked internal stop codons. Using the standard codon table, each CDS was translated into its corresponding protein sequence. Redundant sequences encoding identical protein sequences were removed. The codon similarity index (CSI) was then calculated for the remaining sequences, and the top 10% with the highest CSI values were retained for subsequent analysis. CSI was calculated as follows.

For each amino acid, the relative adaptiveness *w*_*ij*_ of codon *i* encoding amino acid *j* is defined as the ratio of the frequency of that codon in the genome (*x*_*ij*_) to the frequency of the most frequently used synonymous codon for that amino acid (*x*_*jmax*_):

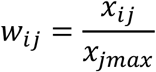

For a CDS containing *L* codons, the CSI is defined as the geometric mean of the relative adaptiveness values across all codons:

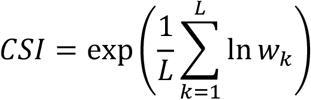

Among the CDSs ranked within the top 10% CSI values, 90% were randomly selected for training and validation, and the remaining 10% were reserved for testing.

To train CodonEXP, we used protein sequences and corresponding in vivo protein abundance data for the same 15 plant species from the MPB-EXP dataset^28^, which was constructed based on the PaxDb database. Protein abundance values were categorized as high abundance (top 33%, labeled as 1) or low abundance (bottom 33%, labeled as 0). To map protein abundance measurements to CDS sequences, we first generated a non-redundant CDS dataset and corresponding translated protein sequences from the reference genomes of the same plant species using the procedure described above for CodonNAT. Protein sequences from PaxDb were then mapped to this non-redundant CDS dataset. Exact sequence matches were accepted directly. If no exact match was found, BLASTp was used to identify similar sequences. From the BLASTp results, sequences with at least 90% sequence similarity and at least 50% alignment coverage were selected, prioritizing matches with the lowest E-value.

This procedure established correspondences between CDS sequences and their associated in vivo protein abundance labels for each species. Redundant sequences were then removed using CD-HIT with a 90% protein sequence identity threshold. The remaining data were divided into training/validation (80%) and testing (20%) sets. Stratified sampling was used to maintain balanced class distributions.

### Model architectures of CodonNAT and CodonEXP

CodonNAT and CodonEXP share a similar model architecture. Protein sequences and CDS sequences are provided as inputs to the ESM2 (650M) and mRNA-FM models, respectively. In mRNA-FM, tokens are defined at the codon level, which corresponds one-to-one with the amino acid tokens used by ESM2 (650M). Because both models use a hidden size of 1280, their output vectors can be combined through weighted addition. The weights used for this combination are trainable parameters optimized during model training. The maximum positional embeddings supported by ESM2 (650M) and mRNA-FM are 1026 and 1024, respectively. To ensure compatibility between the two models during vector addition, the maximum sequence length in our model was set to 1024 positions. Sequences shorter than this length were padded using the <pad> token.

As a result, each input protein−CDS pair is represented as a matrix with dimensions (1024, 1280), where each aligned amino acid−codon pair corresponds to a 1280-dimensional feature vector. In the CodonNAT model, the vector at each position is used directly to predict the codon category at that position. In contrast, the CodonEXP model applies an attention mechanism to aggregate all amino acid−codon pair vectors into a single 1280-dimensional vector representing the entire sequence. This vector is then passed through fully connected layers to predict the probability that the input sequence corresponds to high protein abundance.

### Training of CodonNAT and CodonEXP models

The CodonNAT model was trained using a masked language modeling task. Under the constraint that the amino acid at each position remains unchanged, each codon in the CDS had a 15% probability of being selected for masking. Among the selected codons, 80% were replaced with the special token <mask>, 10% were replaced with a randomly selected codon, and the remaining 10% were left unchanged. The model was trained to predict the original codon category at the masked positions. The dataset was divided into training and validation sets with a ratio of 9:1. The learning rate was linearly decayed from 1e^−4^ to zero during training. Training was performed for up to 50 epochs with a batch size of 2 on a single H20 GPU. Early stopping was applied if prediction accuracy on masked positions in the validation set did not improve for five consecutive epochs. The training loss was computed using a multi-class cross-entropy loss calculated only at the masked positions. Given the set of masked positions *M*, the true labels *y*_*i*_, and the model-predicted probability distribution *p*_*θ*_, the loss function *ℒ* is defined as:

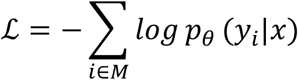

For CodonEXP training, a five-fold cross-validation strategy was used. The learning rates for the linear layers and the pre-trained language model layers were linearly decayed from 1e^−3^ and 1e^−5^ to zero, respectively. Training was performed with a batch size of 2 on a single H20 GPU for 20 epochs per fold. For each fold, the model with the highest F1 score on the validation set was retained. Training used a joint supervision strategy consisting of a primary prediction branch and an auxiliary prediction branch. The primary branch receives integrated features derived from both CDS and protein sequences, whereas the auxiliary branch uses only CDS-derived features. Both branches output raw logits corresponding to the positive class in the binary classification task. The final loss is defined as the sum of the losses from both branches. Both branches use the PyTorch binary cross-entropy loss with logits (BCEWithLogitsLoss):

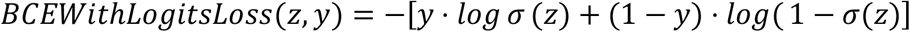

where *z* represents the raw logits predicted by the model, *y* is the binary label (0 or 1), and *σ* is the sigmoid function defined as

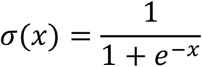

The overall CodonEXP loss function is therefore

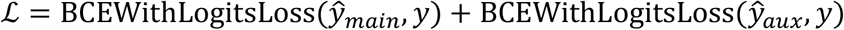

where *ŷ*_*main*_ and *ŷ*_*aux*_ are the unactivated logits produced by the primary and auxiliary branches, respectively.

### Cross-species prediction accuracy of BFC, CodonNAT, and CodonEXP

After removing test sequences showing more than 90% similarity to any training sequence using CD-HIT, prediction accuracies were calculated for each species-specific test set using BFC, CodonNAT, and CodonEXP models trained on datasets from 15 different species. Based on these accuracy values, similarities among species in terms of prediction performance were measured using correlation distance, defined as one minus the Pearson correlation coefficient. Hierarchical clustering with complete linkage was then performed to generate a species clustering tree. For visualization purposes, the node heights in the clustering tree were linearly rescaled into an arithmetic sequence, producing evenly distributed branch lengths while preserving the original tree topology and clustering relationships. The resulting clustering order was used to arrange rows in the heatmap so that species with similar prediction performance appear adjacent to one another.

### Normalization of accuracy differences and codon diversity quantification

To remove species-specific effects when comparing prediction accuracy differences between CodonNAT and BFC across amino acids (AAs), Min−Max normalization was applied within each species. For each species *s* and amino acid *a*, the prediction accuracy difference was calculated as

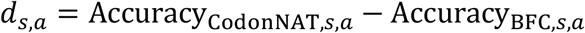

These differences were then normalized within each species by

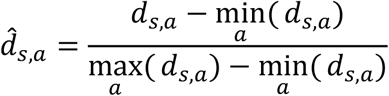

The mean normalized accuracy difference for each amino acid was then calculated across all species

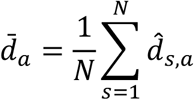

where *N* is the number of species.

To examine the relationship between prediction improvement and codon diversity, codon diversity for each amino acid was quantified using Shannon entropy

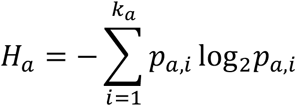

where *k*_*a*_ represents the number of synonymous codons encoding amino acid *a*, and *p*_*a,i*_ denotes the frequency of the *i*^th^ codon.

### Fitness prediction and correlation analysis for the *ccdA* gene

We performed fitness prediction using the benchmark gene *ccdA* from *Escherichia coli*, which contains 62 synonymous mutations with experimentally measured fitness values. To evaluate the ability of CodonNAT to capture the effects of synonymous codon substitutions on fitness, the wild-type CDS and its corresponding amino acid sequence were provided as input to the trained CodonNAT model. For each synonymous mutation, the log-likelihood ratio of the mutant codon relative to the wild-type codon at the mutated position was calculated as

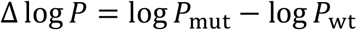

where *P*_mut_ and *P*_wt_ denote the CodonNAT-predicted probabilities of the mutant and wild-type codons, respectively. For the BFC method, the predicted fitness score was calculated as the logarithm of the ratio between the usage frequencies of the mutant and wild-type codons derived from codon frequency tables

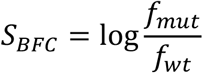

where *f*_*mut*_ and *f*_w*t*_ represent the codon usage frequencies of the mutant and wild-type codons. Spearman’s rank correlation coefficients were then calculated between the predicted scores (Δlog *P* for CodonNAT and *S*_*BFC*_ for BFC) and the experimentally measured fitness values to evaluate model performance. The previously reported Spearman correlation coefficient for CodonTransformer was used for comparison.

### Procedure of CodonIni, CodonGa, and CodonHa

In the codon optimization workflow, the CodonNAT module is first used to generate an initial CDS from the input protein sequence. Specifically, an initial CDS template is created with the same length as the protein sequence, in which every codon position is filled with the <mask> token. CodonNAT then predicts, for each position, the synonymous codon corresponding to the amino acid at that site with the highest probability. The resulting sequence forms the complete initial CDS. This procedure is referred to as CodonIni.

The initial CDS is subsequently optimized using either a genetic algorithm (CodonGa) or hallucination-based design (CodonHa). Optimization is guided by a Fitness score *F*,as the product of the Naturalness score *N* predicted by CodonNAT and the Probability of high abundance *E* predicted by CodonEXP

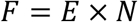

The Fitness score directs the optimization process toward the final CDS.

The iterative optimization of the initial CDS using a genetic algorithm is referred to as CodonGa. The algorithm begins by generating an initial population of sequence variants from the input CDS, introducing synonymous codon substitutions at a rate of 5%. Each individual in the population therefore represents a unique combination of synonymous codons. During evaluation, the algorithm uses five independent models from CodonEXP to estimate the probability of high protein abundance

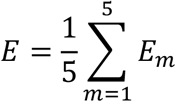

where *E*_*m*_ represents the predicted probability of high expression from the *m*^*th*^ model. At the same time, sequence Naturalness is estimated through repeated stochastic sampling. For each sampling step, each codon position has a 15% probability of being selected. Among the selected codons, 80% are replaced with the <mask> token, 10% are substituted with a random codon, and the remaining 10% remain unchanged. The Naturalness score N for a sequence is calculated as

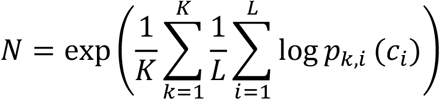

where *K* = 10 represents the number of randomized samplings, *L* denotes the number of sampled codons per sampling, and *p*_*k,i*_(*c*_*i*_) is the CodonNAT-predicted probability of codon *c*_*i*_ at position *i* during the *k*^*th*^ sampling. During each generation, the sequence with the highest Fitness score is preserved in an elite pool and carried forward unchanged to the next generation. In addition, the top 20% of sequences ranked by *F* are selected for crossover and mutation. During crossover, a random codon position is selected as the cut point between two sequences, and the offspring sequence inherits the segment before the cut point from one parent and the segment after the cut point from the other. Mutation introduces synonymous codon substitutions at 5% of codon positions in each individual. After generating the new population, the algorithm recalculates *E, N*, and *F* for all sequences. The optimization process terminates when the maximum Fitness score does not improve for 10 consecutive generations or when the number of iterations reaches 100. For optimization of mCry2Ab and TNFα, a population size of 200 was used; for other proteins, a population size of 100 was applied.

The hallucination-based optimization procedure is referred to as CodonHa. After generating the initial CDS, the algorithm performs gradient-guided iterative optimization of the sequence. During each iteration, the algorithm balances improvements in predicted expression probability *E* with preservation of codon context Naturalness *N*.

For the current sequence, each CodonEXP model obtained through five-fold cross-validation computes a gradient vector ∇_*i*_ at the embedding layer for each codon position *i*. This vector represents the direction and magnitude of change in predicted expression probability resulting from small perturbations of the codon embedding. For the *i*^*th*^ codon *c*_*i*_, with embedding *E*(*c*_*i*_) and a candidate synonymous codon *c*′_*i*_ with embedding *E*(*c*′_*i*_), the contribution of this substitution to predicted expression improvement according to the *m*^*th*^ model is

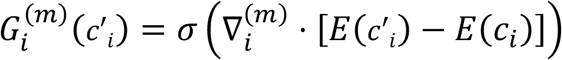

where *σ* is the sigmoid function.

The contributions from the five models are averaged to obtain the overall expression gain for substitution *c*′_*i*_

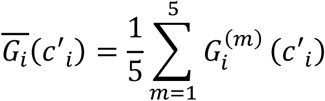

To maintain the optimized CDS within the natural codon usage range of the host species, CodonHa incorporates additional constraints based on CodonNAT-predicted probabilities and synonymous codon usage distributions. This is implemented using a Jensen−Shannon divergence (JSD)−based regularization term. For each amino acid, the synonymous codon usage frequencies in the current CDS define a distribution *P*_*aa*_. The reference codon usage distribution for the host genome is denoted as *Q*_*aa*_. The divergence between the two distributions is defined as

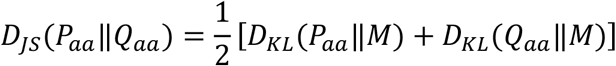

where

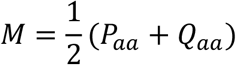

and *D*_*KL*_ denotes the Kullback-Leibler divergence

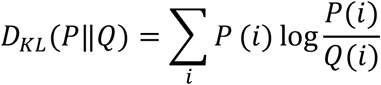

Here *P*(*i*) represents the frequency of synonymous codon i in the current optimized CDS, and *Q*(*i*) represents the reference codon usage frequency in the host genome. When a synonymous substitution *c*′_*i*_ occurs, codon usage frequencies for the corresponding amino acid are recalculated, producing a new divergence *D*′_*IS*_. The improvement in codon usage compatibility is defined as

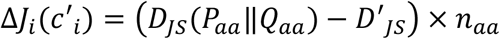

where *n*_*aa*_ represents the number of occurrences of that amino acid in the sequence. This value is normalized using the sigmoid function

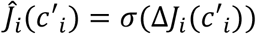

For each synonymous substitution *c*′_*i*_, a combined substitution score is defined as

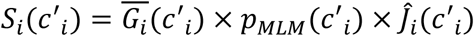

where *p*_*MLM*_(*c*′_*i*_) is the CodonNAT-predicted probability for codon *c*′_*i*_ at that position.

Each iteration consists of two stages. In the first stage, candidate substitutions are restricted to those for which both the expression gain 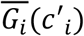 and the JSD gain *ĵ*_*i*_ (*c*′_*i*_) exceeds 0.5. For each position, the substitution with the highest score *S*_*i*_(*c*′_*i*_) is identified. The top 15% of substitutions ranked *S*_*i*_ across all positions are then applied to generate an updated CDS.

In the second stage, the JSD constraint is relaxed. Only the expression gain must exceed a threshold defined as min(0.5,1 − *E*). Again, the substitution with the highest *S*_*i*_(*c*′_*i*_) is selected for each position, and the top 15% of substitutions are applied to produce the optimized CDS for that iteration.

Starting from the initial CDS generated by CodonIni, this iterative optimization process is repeated. After each iteration, the values of *F, E*, and *N* are recalculated. Optimization stops when either the total number of iterations reaches 96 or when *F* fails to improve for 20 consecutive iterations while both *E* and *N* exceed predefined thresholds (0.9 and 0.6). The CDS with the highest Fitness score encountered during the optimization process is selected as the final optimized sequence.

In the Ha-GC3 variant, the substitution score *S*_*i*_(*c*′_*i*_) for a candidate synonymous codon *c*′_*i*_ at position *i* is modified by introducing a GC3 bias term

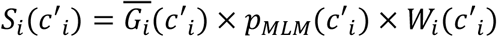

where the GC3 weighting term *W*_*i*_(*c*′_*i*_) is defined as

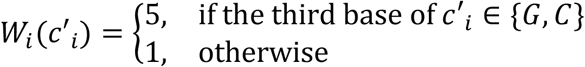

This weighting encourages substitutions that introduce G or C at the third codon position during optimization.

### Additional codon optimization methods

Several existing codon optimization methods were evaluated.The BFC method selects the most frequently used codon for each AA based on species-specific codon usage tables. Genewiz (https://www.genewiz.com/public/services/gene-synthesis/codon-optimization) provides a commercial codon optimization service. CodonTransformer (https://adibvafa.github.io/CodonTransformer/GoogleColab) uses a Transformer-based neural network to optimize codon sequences by modeling context-dependent codon usage. GC3 Recoder (https://larrywu.shinyapps.io/GC3-recoder) increases GC content at the third codon position to potentially enhance gene expression.

### Metrics for evaluating optimized codons

Multiple metrics were used to evaluate codon optimization results generated by different tools, including the Jaccard index, sequence similarity, CAI, and GC3 content.

The Jaccard index is a set-based similarity measure defined as the ratio of the size of the intersection to the size of the union of two sets. It quantifies the proportion of shared elements between two groups. For two CDS sequences *X* and *Y*, with corresponding codon sets *A* and *B*, the Jaccard index is defined as

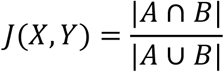

This index ranges from 0 to 1, where higher values indicate greater similarity between the codon sets of the two sequences and therefore more consistent codon composition.

Sequence similarity measures the proportion of matching codons at corresponding positions between two sequences of equal length. For two sequences *A* and *B* of length *L*, sequence similarity is calculated as

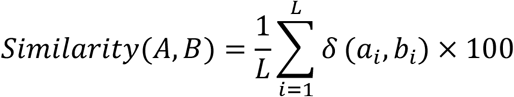

where *a*_*i*_ and *b*_*i*_ denote the codons at position *i* in sequences *A* and *B*, respectively. The indicator function *δ*(*a*_*i*_, *b*_*i*_) equals 1 when *a*_*i*_ = *b*_*i*_ and 0 otherwise.

The CAI calculation is conceptually similar to CSI but uses a set of highly expressed reference genes to compute the relative adaptiveness of each codon. CAI values for each sequence were calculated using the GenScript codon analysis tool (https://www.genscript.com/tools/rare-codon-analysis)^47^.

The GC3 content of a CDS is defined as the proportion of guanine (G) and cytosine (C) nucleotides occurring at the third position of codons in the sequence. If a CDS contains *N* codons and *b*_3*i*_ denotes the nucleotide at the third position of the *i*^*th*^ codon, then

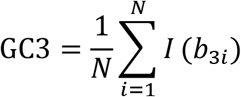

where the indicator function *I*(*b*_3*i*_) is defined as

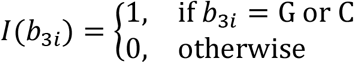

### DNA sequence synthesis and plasmid construction

The CDS sequences optimized by different codon optimization methods were synthesized by Suzhou Junji Gene Technology Co., Ltd. All synthesized CDS sequences were fused to a C-terminal 6×His tag for detection of protein expression. The synthesized target sequences were cloned into the plant binary expression vector pEarleyGate104, positioned between the 35S promoter and the OCS terminator, using the ClonExpress Ultra One Step Cloning Kit (C115-01, Vazyme Biotech Co., Ltd). All plasmid construction and amplification steps were performed in Escherichia coli strain DH5α (DL1001, Shanghai Weidi Biotechnology Co., Ltd). The 18 expected constructs were verified by Sanger sequencing and subsequently transformed into Agrobacterium tumefaciens strain GV3101 (AC1001, Shanghai Weidi Biotechnology Co., Ltd) for further experiments.

### Protein expression and western blot analysis

To ensure consistent expression conditions and minimize systematic variation, Agrobacterium suspensions carrying six different codon-optimized constructs were adjusted to the same infiltration concentration (OD600 = 0.5) and injected in equal volumes into different regions of the same tobacco leaf for transient expression. Seventy-two hours after infiltration, two leaf discs (10 mm in diameter) were collected from each injection site to ensure uniform sampling area and biomass. Samples were ground into powder, after which 2× SDS loading buffer was added. The mixture was incubated on ice for 10 minutes and then denatured at 100°C for 10 minutes. After cooling, samples were centrifuged at 12,000 rpm for 3 minutes, and the supernatant was collected as the total protein extract. Protein samples were separated using 10% SDS−PAGE and then wet-transferred onto a 0.45 μm PVDF membrane. After transfer, the membrane was stained with Ponceau S and rinsed with TBST to remove residual dye. The PVDF membrane was then blocked with 5% skim milk at room temperature for 1 hour. After washing with TBST, the membrane was incubated overnight at 4°C with anti-His-Tag monoclonal primary antibody (diluted 1:10,000 in 3% BSA solution; AE003, ABclonal). Following primary antibody incubation and washing, the membrane was incubated with HRP-conjugated goat anti-mouse IgG (H+L) secondary antibody (diluted 1:5,000; AS003, ABclonal) at room temperature for 1 hour. After removal of unbound secondary antibody, chemiluminescent detection was performed using the Clarity Western ECL substrate kit (1705060; Bio-Rad), and signals were captured using a Tanon-4600 imaging system. For quantitative analysis of protein expression, band intensities in the chemiluminescent images were measured using ImageJ software. The RuBisCO large subunit band (~55 kDa), visualized by Ponceau S staining, was used as an internal reference for normalization and statistical analysis of the target protein bands.

### Fluorescence Imaging

Fluorescence imaging was performed at the Public Instrument Platform of the College of Agronomy and Biotechnology, China Agricultural University, using a Zeiss LSM880 inverted confocal microscope. Excitation was provided by a HeNe red laser (633 nm) and a 561 nm laser, with laser power maintained between 0.6% and 2.4%. For each sample, six random fields of view were imaged using a 10× objective lens (NA 0.45). Laser intensity was kept constant throughout imaging to ensure comparability between samples. Red fluorescence intensity in the captured images was quantified using ImageJ software.

## Additional information

### Data availability

The datasets used for training and testing CodonNAT and CodonEXP are available at https://zenodo.org/records/19199736. Data corresponding to Figs. 2a, 2b, 2d, 2e, 3a, 3b, 3c, 3d, 3e, 3f, 4a, 4b, 4c, 4d, 4e, 4f, 5b, 5h and Supplementary Figs. 1, 2, 3, 4, 6, 7 are available as Source Data files. All source data are available with this paper.

### Code availability

The source codes and a user-friendly web interface of HalluCodon are available at https://github.com/YuxuanLou/HalluCodon and https://codon.oneshot.ac.cn, respectively. We have trained the CodonNAT and CodonEXP models separately on 15 plant species. The species names and their corresponding weight storage paths are deposited in https://zenodo.org as follows accessions: Arabidopsis (19126265), Canola (19129186), Sweet orange (19133772), Cotton (19135167), Soybean (/19135889), Barley (19136614), Medicago (19143273), Tobacco (19143653), Rice (19144006), Earthmoss (19144435), Tomato (19150439), Potato (19150918), Wheat (19151412), Grape (19151918), Maize (19152348).

## Acknowledgements

This research was supported by the Fundamental and Interdisciplinary Disciplines Breakthrough Plan of the Ministry of Education of China (JYB2025XDXM705), the Guiding Special Fund for Central Universities to Build World-Class Universities (Disciplines) and Promote Characteristic Development (2025AC030), and the Pinduoduo-China Agricultural University Research Fund (PC2024A02002).

## Author contributions

X.W. and Y.X.L. conceived the project. X.W. and J.Y. designed and supervised the study. Y.X.L., F.X., and Z.Z. performed the bioinformatic analyses and model development. S.M., Y.L., and Y.T. performed experimental validation. Q.C. and T.W. developed the web interface. Y.X.L., J.Y., and X.W. wrote the manuscript. All authors reviewed and edited the manuscript and contributed intellectual input.

## Tables and Figures

**Supplementary Table 1.** Data from the fifteen plant species for training base model.

| Latin name of host species<br>(assembly version) | Common<br>name | PaxDb dataset<br>(protein abundance<br>dataset) | Sample<br>size for<br>CodonNAT | Sample size<br>for<br>CodonEXP |
| --- | --- | --- | --- | --- |
| *Arabidopsis thaliana*<br>(TAIR10.1) | Arabidopsis | 3702-WHOLE_ORGANISM<br>-integrated | 4402 | 13103 |
| *Brassica napus*<br>(AST_PRJEB5043_v1) | Canola | 3708-PXD005787_Geng_Fr<br>ont_Mol_Biosci_2017<br>_Bnapus_guard_cell | 10587 | 2949 |
| *Citrus sinensis* (DVS_A1.0) | Sweet orange | 2711-PXD008366_leaf | 3539 | 1788 |
| *Gossypium hirsutum*<br>(Gossypium_hirsutum_v2.1) | Cotton | 3635-LEAF-integrated | 9425 | 4245 |
| *Glycine max*<br>(Glycine_max_v2.1) | Soybean | 3847-SHOOT-integrated | 7993 | 1512 |
| *Hordeum vulgare* (MorexV3) | Barley | 4513-ENDOSPERM-integrated | 3914 | 1705 |
| *Medicago truncatula*<br>(MedtrA17_4.0) | Medicago | 3880-PXD002692_flower | 6188 | 7385 |
| *Nicotiana tabacum*<br>(ASM71507v2) | Tobacco | 4097-LEAF-integrated | 7720 | 3112 |
| *Oryza sativa* (IRGSP-1.0) | Rice | 39947-WHOLE_ORGANISM<br>-integrated | 4252 | 6169 |
| *Physcomitrella patens*<br>(Phypa V3) | Earthmoss | 3218-PXD013606_Physco<br>mitrella_patens | 5130 | 5652 |
| *Solanum lycopersicum*<br>(SL3.0) | Tomato | 4081-PXD012810_McWhite<br>_Cell_2020_Tomato_<br>green | 3767 | 4106 |
| *Solanum tuberosum*<br>(SoITub_3.0) | Potato | 4113-LEAF-integrated | 5274 | 4302 |
| *Triticum aestivum* (IWGSC<br>CS RefSeq v2.1) | Wheat | 4565-LEAF-integrated | 13438 | 2600 |
| *Vitis vinifera* (PN40024 v4) | Grape | 29760-SHOOT-integrated | 4165 | 1329 |
| *Zea mays* (Zm-B73-<br>REFERENCE-NAM-5.0) | Maize | 4577-LEAF-integrated | 6738 | 6520 |

### Supplementary Dataset

Detailed information, protein sequences, and original and optimized CDSs for the fifteen benchmark proteins used for model evaluation and experimental validation.

**Supplementary Fig. 1.**
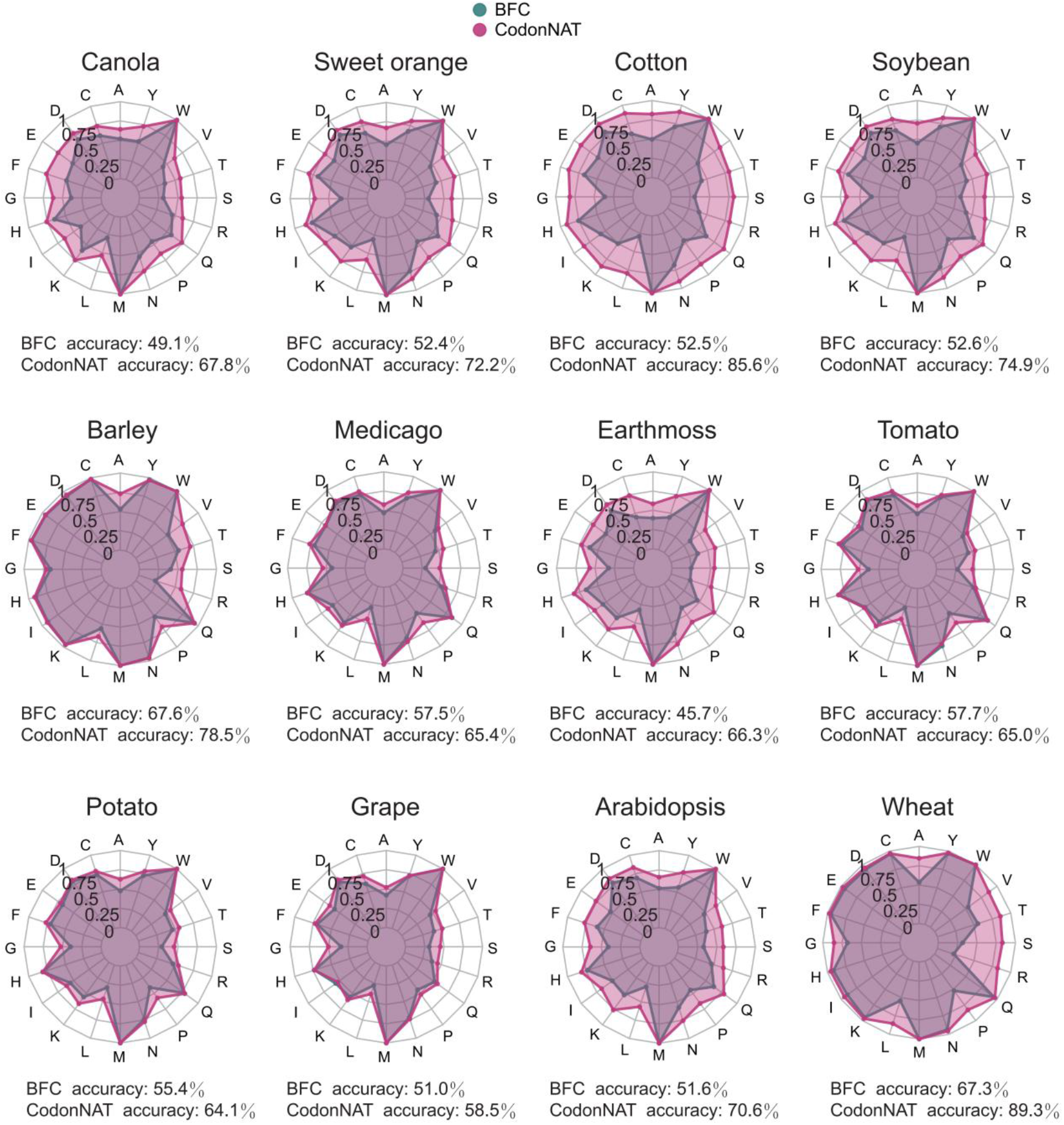
Comparison of synonymous codon prediction accuracy between CodonNAT and the BFC model across fifteen plant species.

**Supplementary Fig. 2.**
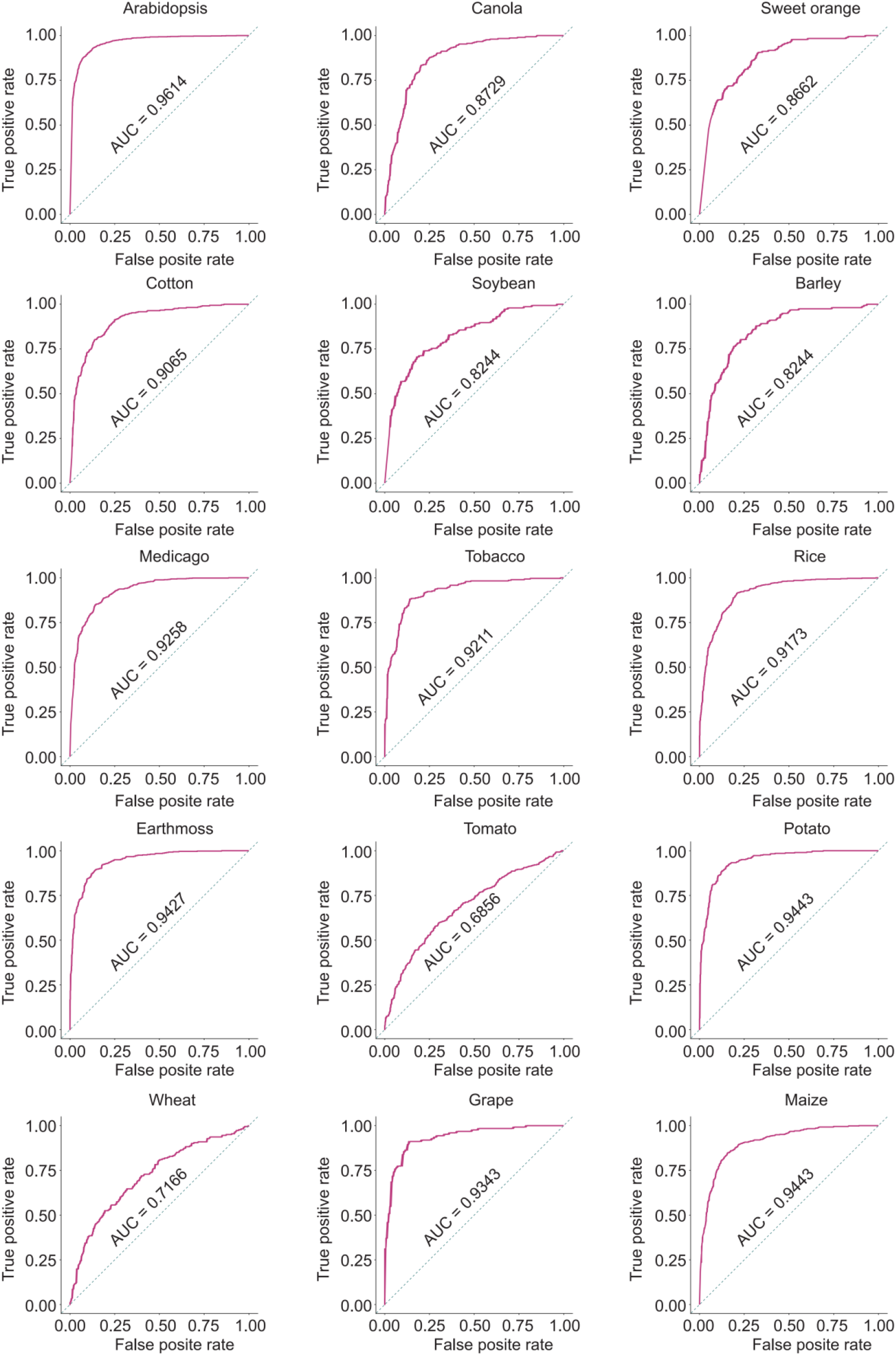
Receiver operating characteristic (ROC) curves of CodonEXP on the testing sets from the fifteen plant species. The probability scores of high protein expression used to generate these curves were obtained by averaging the outputs of the five models trained during five-fold cross-validation.

**Supplementary Fig. 3.**
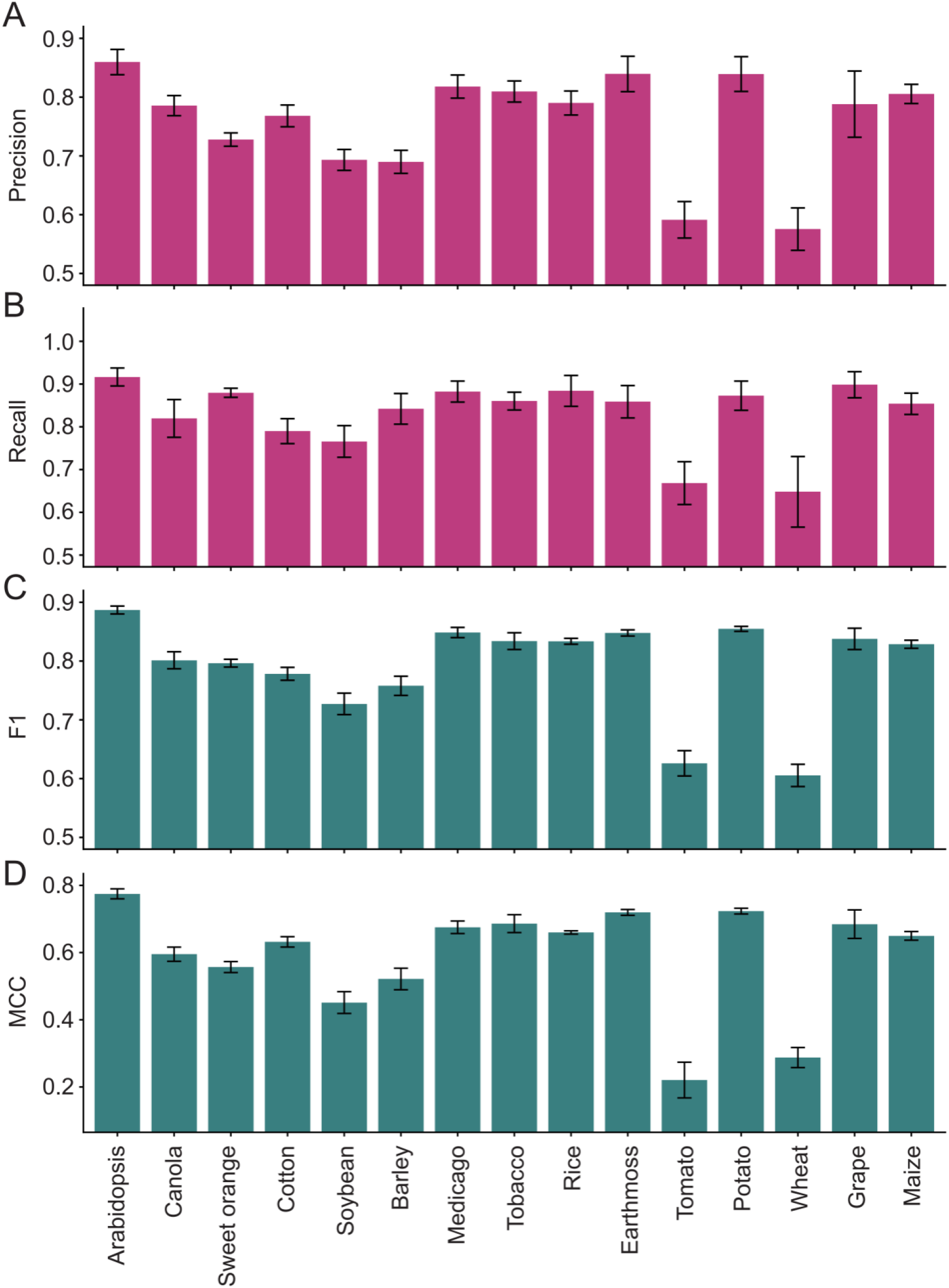
Performance metrics for CodonEXP across the fifteen plant species, including precision (A), recall (B), F1 score (C), and Matthews correlation coefficient (MCC) (D). Error bars represent standard deviations calculated from five-fold cross-validation.

**Supplementary Fig. 4.**
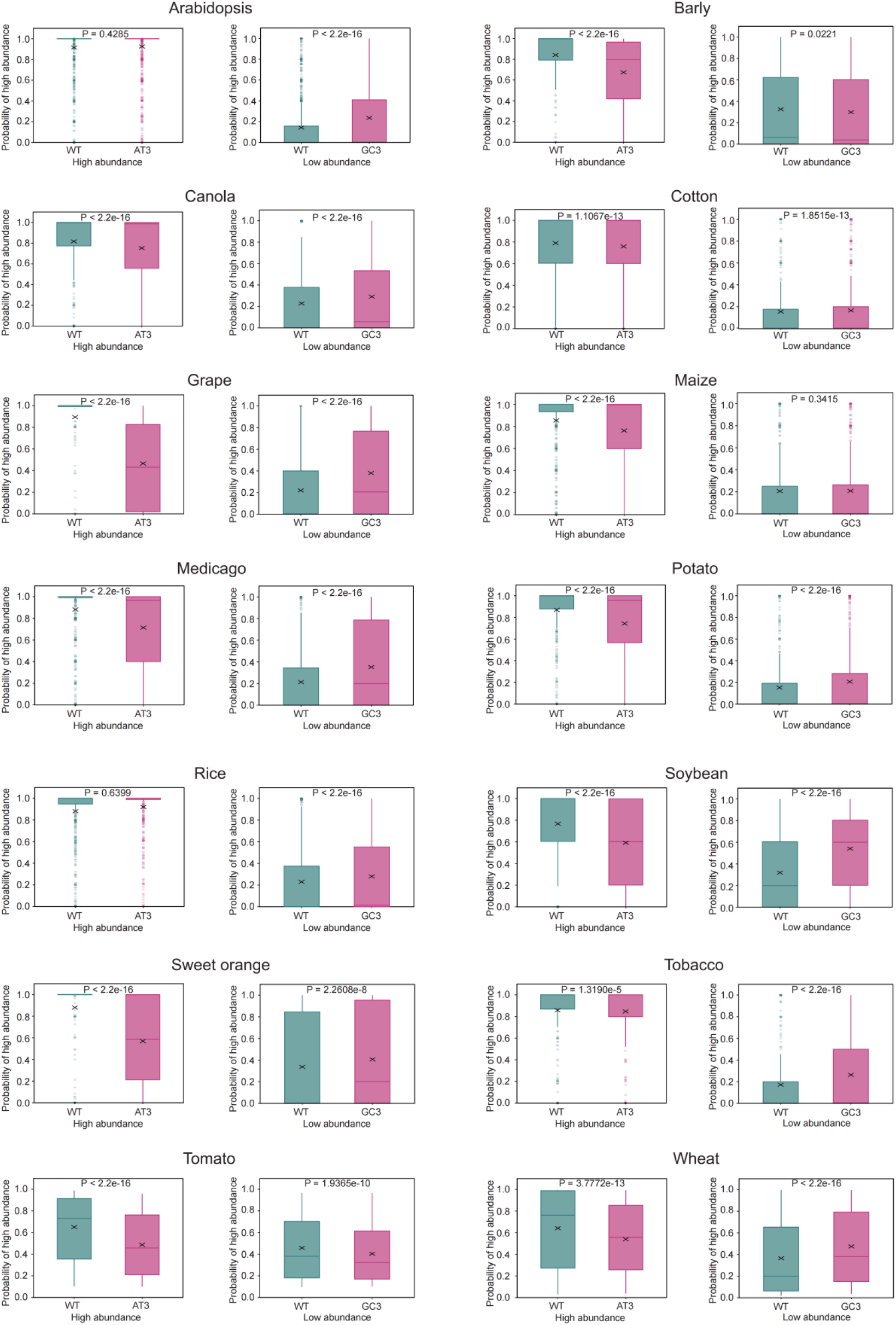
Comparison of predicted probabilities of high abundance by CodonEXP between original CDSs and synonymous codon variants with altered GC3 content in fourteen plant species. Statistical significance was assessed by the paired Wilcoxon signed-rank test, with exact p-values indicated above the boxes. Each boxplot displays group mean values (marked with an X) and corresponding predicted probabilities of high abundance.

**Supplementary Fig. 5.**
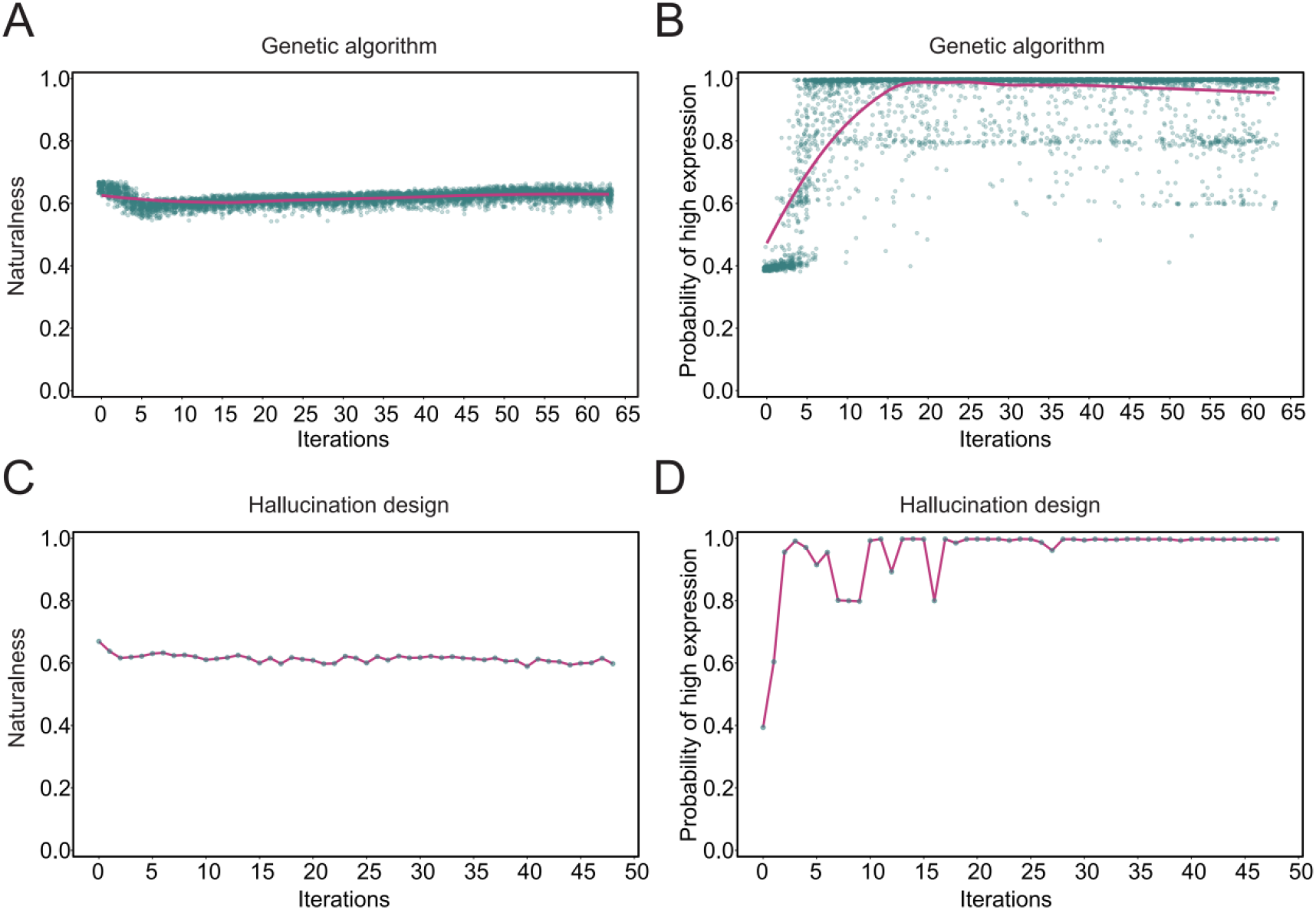
Trends in naturalness scores during iterative optimization of DsRed2 sequences using CodonGa (a) and CodonHa (b). Trends in probability scores of high protein expression during iterative optimization of DsRed2 sequences using CodonGa (c) and CodonHa (d).

**Supplementary Fig. 6.**
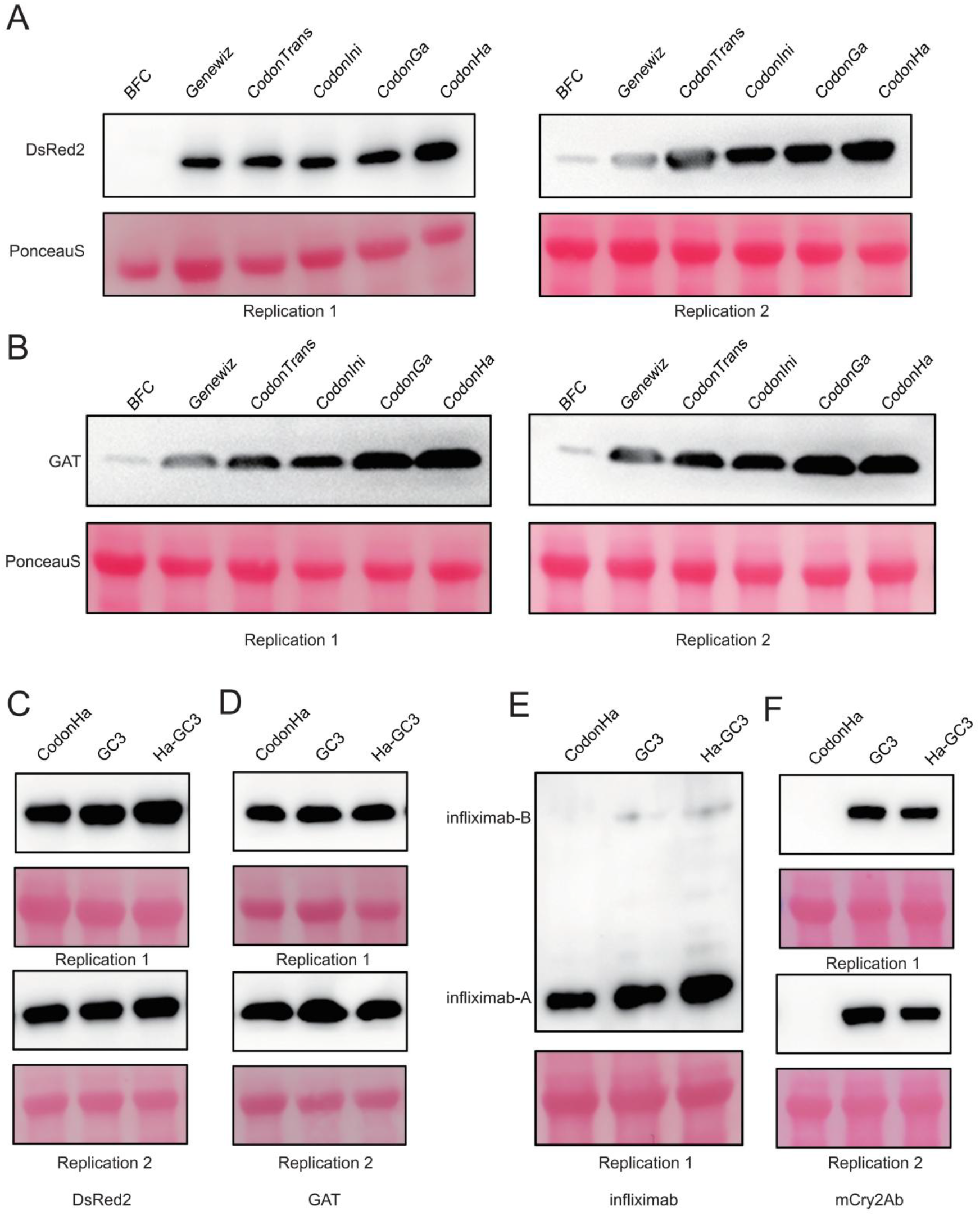
Western blot analysis of protein expression levels for DsRed2 (a) and GAT (b) sequences optimized using five methods. Western blot analysis of protein expression levels for DsRed2 (c), GAT (d), infliximab-AB (e), and mCry2Ab (f) sequences optimized using CodonHa alone, arbitrary GC3 enrichment at 0.8 content (GC3), and GC3 enrichment incorporated with CodonHa (Ha-GC3).

**Supplementary Fig. 7.**
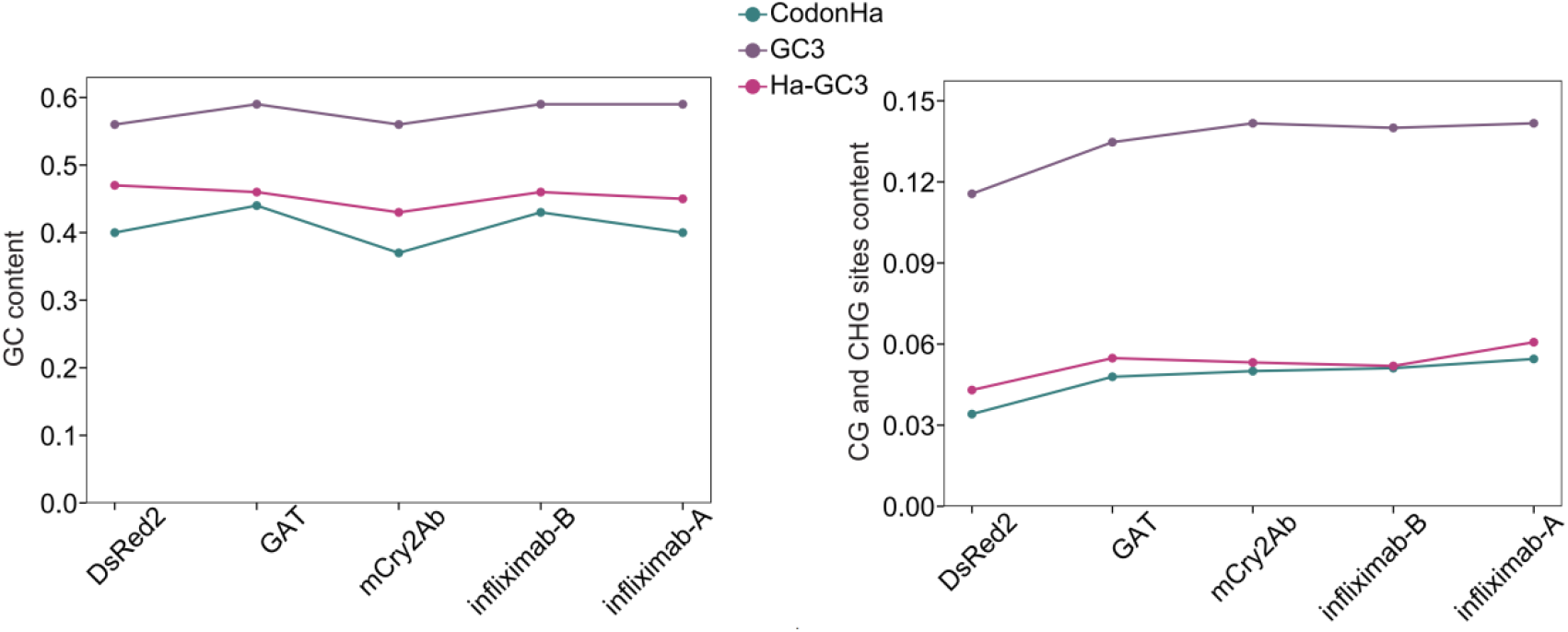
GC content (left panel) and methylation site content (CG and CHG) (right panel) of sequences optimized using CodonHa, GC3, and Ha-GC3.

## References

1. Nirenberg, M.W. & Matthaei, J.H. The dependence of cell-free protein synthesis in E. coli upon naturally occurring or synthetic polyribonucleotides. Proceedings of the National Academy of Sciences 47, 1588–1602 (1961).

2. Bernfield, M.R. & Nirenberg, M.W. RNA Codewords and Protein Synthesis. Science 147, 479–484 (1965).

3. Hershberg, R. & Petrov, D.A. Selection on Codon Bias. Annual Review of Genetics 42, 287–299 (2008).

4. Pechmann, S. & Frydman, J. Evolutionary conservation of codon optimality reveals hidden signatures of cotranslational folding. Nature Structural & Molecular Biology 20, 237–243 (2013).

5. Rocha, E.P.C. Codon usage bias from tRNA’s point of view: Redundancy, specialization, and efficient decoding for translation optimization. Genome Research 14, 2279–2286 (2004).

6. Mauger, D.M. et al. mRNA structure regulates protein expression through changes in functional half-life. Proceedings of the National Academy of Sciences 116, 24075–24083 (2019).

7. Pertzev, A.V. & Nicholson, A.W. Characterization of RNA sequence determinants and antideterminants of processing reactivity for a minimal substrate of Escherichia coli ribonuclease III. Nucleic Acids Research 34, 3708–3721 (2006).

8. Li, G. et al. Integrated biotechnological and AI innovations for crop improvement. Nature 643, 925–937 (2025).

9. Fausther-Bovendo, H. & Kobinger, G. Plant-made vaccines and therapeutics. Science 373, 740–741 (2021).

10. Sharp, P.M. & Li, W.-H. The codon adaptation index-a measure of directional synonymous codon usage bias, and its potential applications. Nucleic Acids Research 15, 1281–1295 (1987).

11. Quax, Tessa E.F., Claassens Nico J., Söll, D. & van der Oost, J. Codon Bias as a Means to Fine-Tune Gene Expression. Molecular Cell 59, 149–161 (2015).

12. Moss, M.J., Chamness, L.M. & Clark, P.L. The Effects of Codon Usage on Protein Structure and Folding. Annual Review of Biophysics 53, 87–108 (2024).

13. Jain, R., Jain, A., Mauro, E., LeShane, K. & Densmore, D. ICOR: improving codon optimization with recurrent neural networks. BMC Bioinformatics 24, 132 (2023).

14. Fallahpour, A., Gureghian, V., Filion, G.J., Lindner, A.B. & Pandi, A. CodonTransformer: a multispecies codon optimizer using context-aware neural networks. Nature Communications 16, 3205 (2025).

15. Han, X. et al. DeepCodon: A deep learning codon-optimization model to enhance protein expression. BioDesign Research 7 (2025).

16. Zhang, H. et al. Deep generative models design mRNA sequences with enhanced translational capacity and stability. Science 390, eadr8470 (2025).

17. Thess, A. et al. Sequence-engineered mRNA Without Chemical Nucleoside Modifications Enables an Effective Protein Therapy in Large Animals. Molecular Therapy 23, 1456–1464 (2015).

18. Karikó, K., Buckstein, M., Ni, H. & Weissman, D. Suppression of RNA Recognition by Toll-like Receptors: The Impact of Nucleoside Modification and the Evolutionary Origin of RNA. Immunity 23, 165–175 (2005).

19. Karikó, K. et al. Incorporation of Pseudouridine Into mRNA Yields Superior Nonimmunogenic Vector With Increased Translational Capacity and Biological Stability. Molecular Therapy 16, 1833–1840 (2008).

20. Govaerts, R., Nic Lughadha, E., Black, N., Turner, R. & Paton, A. The World Checklist of Vascular Plants, a continuously updated resource for exploring global plant diversity. Scientific Data 8, 215 (2021).

21. Xie, L. et al. Technology-enabled great leap in deciphering plant genomes. Nature Plants 10, 551–566 (2024).

22. Schreiber, M., Jayakodi, M., Stein, N. & Mascher, M. Plant pangenomes for crop improvement, biodiversity and evolution. Nature Reviews Genetics 25, 563–577 (2024).

23. Sang, T., Zhang, Z., Liu, G. & Wang, P. Navigating the landscape of plant proteomics. Journal of Integrative Plant Biology 67, 740–761 (2025).

24. Guo, W. et al. A barley pan-transcriptome reveals layers of genotype-dependent transcriptional complexity. Nature Genetics 57, 441–450 (2025).

25. Zhu, W. et al. Large-scale translatome profiling annotates the functional genome and reveals the key role of genic 3′ untranslated regions in translatomic variation in plants. Plant Communications 2, 100181 (2021).

26. Lin, Z. et al. Evolutionary-scale prediction of atomic-level protein structure with a language model. Science 379, 1123–1130 (2023).

27. Shen, T. et al. Accurate RNA 3D structure prediction using a language model-based deep learning approach. Nature Methods 21, 2287–2298 (2024).

28. Liu, T. et al. Effective Gene Expression Prediction and Optimization from Protein Sequences. Advanced Science 12, 2407664 (2025).

29. Vogel, C. et al. Sequence signatures and mRNA concentration can explain two-thirds of protein abundance variation in a human cell line. Molecular Systems Biology 6, MSB201059 (2010).

30. Zur, H. & Tuller, T. Transcript features alone enable accurate prediction and understanding of gene expression in S. cerevisiae. BMC Bioinformatics 14, S1 (2013).

31. Nikolados, E.-M., Wongprommoon, A., Aodha, O.M., Cambray, G. & Oyarzún, D.A. Accuracy and data efficiency in deep learning models of protein expression. Nature Communications 13, 7755 (2022).

32. Li, T. et al. Modeling 0.6 million genes for the rational design of functional cis-regulatory variants and de novo design of cis-regulatory sequences. Proceedings of the National Academy of Sciences 121, e2319811121 (2024).

33. Khare, E. et al. Discovering design principles of collagen molecular stability using a genetic algorithm, deep learning, and experimental validation. Proceedings of the National Academy of Sciences 119, e2209524119 (2022).

34. Anishchenko, I. et al. De novo protein design by deep network hallucination. Nature 600, 547–552 (2021).

35. Pacesa, M. et al. One-shot design of functional protein binders with BindCraft. Nature 646, 483–492 (2025).

36. Wang, M. et al. PaxDb, a Database of Protein Abundance Averages Across All Three Domains of Life*. Molecular & Cellular Proteomics 11, 492–500 (2012).

37. Chandra, S. et al. The High Mutational Sensitivity of ccdA Antitoxin Is Linked to Codon Optimality. Molecular Biology and Evolution 39 (2022).

38. Nieuwkoop, T. et al. Revealing determinants of translation efficiency via whole-gene codon randomization and machine learning. Nucleic Acids Research 51, 2363–2376 (2023).

39. Yu, H. et al. An interpretable RNA foundation model for exploring functional RNA motifs in plants. Nature Machine Intelligence 6, 1616–1625 (2024).

40. Outeiral, C. & Deane, C.M. Codon language embeddings provide strong signals for use in protein engineering. Nature Machine Intelligence 6, 170–179 (2024).

41. Bevis, B.J. & Glick, B.S. Rapidly maturing variants of the Discosoma red fluorescent protein (DsRed). Nature Biotechnology 20, 83–87 (2002).

42. Kaufman, I.D., Wu, H.-Y.L. & Hsu, P.Y. GC3 codons enhance protein production in diverse GC- and AT-rich plant species. bioRxiv, 2025.2011.2025.690434 (2025).

43. Real, R. & Vargas, J.M. The Probabilistic Basis of Jaccard’s Index of Similarity. Systematic Biology 45, 380–385 (1996).

44. Kosuri, S. & Church, G.M. Large-scale de novo DNA synthesis: technologies and applications. Nature Methods 11, 499–507 (2014).

45. Sahdev, S., Saini, S., Tiwari, P., Saxena, S. & Singh Saini, K. Amplification of GC-rich genes by following a combination strategy of primer design, enhancers and modified PCR cycle conditions. Molecular and Cellular Probes 21, 303–307 (2007).

46. Zan, Y. et al. The genome and GeneBank genomics of allotetraploid Nicotiana tabacum provide insights into genome evolution and complex trait regulation. Nature Genetics 57, 986–996 (2025).

47. Fan, K., Li, Y., Chen, Z. & Fan, L. GenRCA: a user-friendly rare codon analysis tool for comprehensive evaluation of codon usage preferences based on coding sequences in genomes. BMC Bioinformatics 25, 309 (2024).

